# Conjugative serine recombinase engineering enables genome integration in transformation-resistant bacteria and bioproduction pathway prototyping

**DOI:** 10.64898/2026.04.17.717921

**Authors:** Michael S. Guzman, Cholpisit Kiattisewee, Amanda M. Robert, Jackson Comes, Ryan A.L. Cardiff, Allan Scott, Margaret Cook, Diego Alba Burbano, Stella Anastasakis, Sarah Grube, Kieran Heiberg, Brian Darst, Daniel Howell, Gara N. Dexter, Robert G. Egbert, Jesse G. Zalatan, Adam M. Guss, Joshua R. Elmore, Alexander S. Beliaev, James M. Carothers

## Abstract

Non-model bacteria offer unique metabolic capabilities for sustainable bioproduction, yet their limited genetic accessibility hinders systematic strain development. Here we present conjugation-based serine recombinase-assisted genome engineering (cSAGE), a portable platform for sequential, site-specific genomic integration in transformation-resistant bacteria. Using *Rhodobacter sphaeroides* as a testbed, we show that genome integration fails by electroporation and is not rescued by disruption of three candidate restriction endonucleases. Conjugative delivery of the same integration vectors recovers genomic integrants at frequencies of approximately 10^−4^ per recipient cell, an improvement of at least five orders of magnitude over electroporation. We demonstrate site-specific genomic integration across eight bacterial hosts spanning four classes and two phyla, as well as three sequential payload integrations using a standardized, non-replicating plasmid toolkit. We then use cSAGE to install the same bioproduction pathway in two *Rhodopseudomonas palustris* strains: the reference strain CGA009 and the environmental isolate P4, for which genetic engineering has not previously been reported. Under anoxygenic photosynthetic growth conditions, engineered P4 produces *p*-vinylphenol from *p*-coumarate, whereas CGA009 engineered with the same construct, does not. By enabling standardized genome engineering and cross-host evaluation of bioproduction pathways, cSAGE provides a general framework for non-model strain prototyping and biotransformation discovery.

## Introduction

Efficient microbial conversion of underutilized waste-derived feedstocks into valuable products is central to a sustainable bioeconomy^1^. Across agricultural, chemical, and wastewater management sectors, large reservoirs of carbon exist in the form of volatile fatty acids^2^, lignocellulosic residues^3^, and industrial off-gases^4^. Harnessing these resources relies on leveraging the native metabolic capabilities of non-model bacteria and environmental isolates, together with engineered functions for efficient chemical and fuel production. Among these, purple nonsulfur bacteria (PNSB) represent a particularly versatile and underexploited group of microorganisms for bioproduction^5^. PNSB couple CO_2_ fixation, photosynthesis, and carbon-efficient aromatic compound degradation within a single organism, allowing them to use low-energy infrared light to catabolize waste carbon feedstocks, including lignin-derived aromatics and syngas^6–10^. Despite their metabolic flexibility and the availability of curated genome-scale metabolic models predicting production capabilities^11,12^, the use of PNSB for sustainable biomanufacturing remains limited by the lack of robust, stable, and portable genetic tools^6,7,13^. Like other non-model organisms, many PNSB strains remain recalcitrant to transformation, creating a major bottleneck for strain engineering^14^.

Over the past decade, many advanced tools have been developed to address barriers in microbial genetics and engineering. However, these tools require tradeoffs among DNA delivery, host compatibility, and genome-integration throughput. For example, approaches such as conventional allelic exchange^15^, CRISPR-Cas systems^16^, transposon-based methods^17^, and recombineering^18^ are often limited by transformation efficiency, host-range compatibility, or the need for host-specific genetic components^16^. These approaches can require host-compatible delivery vectors, replication origins, expression systems, selectable markers, or counterselection strategies. Methods that rely on replicating plasmids also require compatible replication origins and may necessitate additional plasmid-curing steps^16–20^. Moreover, replicating plasmids typically require prior screening of functional genetic parts for each new organism^21^. As a result, genome integration strategies that perform well in some species often fail to generalize across bacteria^22^.

Beyond transformation efficiency and host range, genome integration throughput presents a further limitation for non-model strain construction^15^. Widely used Tn7 transposon-based strategies typically support a single integration event^19,23^ at the conserved *attTn7* site, after which target immunity limits reuse of that site. CRISPR-Cas systems require cloning of strain-specific homology-directed repair plasmids for each successive modification^20^. Although both approaches can be made compatible with conjugative DNA delivery^19,23^ successive modifications require either additional genomic target sites or newly designed repair constructs. Recombinase-based landing-pad systems such as CRAGE similarly support conjugative delivery and broad cross-phylum host range, but current implementations support at most two sequential integration sites^24,25^. To our knowledge, no current platform combines conjugative delivery, multiple orthogonal integration sites, marker recycling, and a standardized non-replicating plasmid toolkit for sequential payload integration across diverse bacterial hosts. This gap limits scalable strain construction in non-model organisms.

Serine recombinase-based integration systems have recently emerged as versatile tools for genome engineering in non-model bacteria^26^. Serine-recombinase-assisted genome engineering (SAGE) enables site-specific, unidirectional integration of DNA into the host chromosome by catalyzing recombination between a plasmid-encoded, phage-derived attachment site (*attP*) and a corresponding bacterial chromosomal attachment site (*attB*)^27^. Once a poly-*attB* landing pad composed of multiple orthogonal *attB* sites is installed in the genome, SAGE supports efficient and stable integration of large or multi-gene payloads by using distinct recombinases to direct successive *attP*-flanked cargos to cognate *attB* sites. This architecture enables sequential payload integration without constructing new host-specific homology arms for each insertion. However, broader use of SAGE has been limited by its reliance on electroporation for DNA delivery. Although electroporation is effective in many bacteria, its performance is strongly host dependent, requiring optimization of electroporation conditions and competent-cell preparation for each new species^15^. Even after extensive optimization, some non-model bacteria remain refractory to electroporation^28^.

Conjugation provides several advantages that make it an attractive alternative to electroporation for DNA delivery. First, conjugation is a highly conserved mechanism of horizontal gene transfer across the bacterial kingdom that functions in hosts ranging from laboratory *E. coli* strains to non-model microbes^29–31^. Second, conjugative DNA is initially transferred as single-stranded DNA and then converted to double-stranded DNA in the recipient cell, which can reduce its exposure to some host restriction barriers relative to delivery of double-stranded DNA^32–34^. Accordingly, several recent broad-host-range genetic toolkits rely on conjugation as a robust and portable DNA-delivery strategy^19,23,25^. Conjugation has become an established method for introducing DNA into bacteria that are otherwise refractory to transformation, including PNSB, gut microbiota, marine and soil bacteria, and extremophiles^29^. We therefore reasoned that coupling conjugative delivery to SAGE could reduce its dependence on host-specific electroporation while preserving sequential, site-specific payload integration.

Here we present conjugation-based SAGE (cSAGE), a portable genome engineering toolkit that combines conjugative delivery with sequential, site-specific payload integration. In cSAGE, all plasmids are designed in the Standard European Vector Architecture (SEVA) format^35,36^ with an RK2/RP4 origin of transfer (*oriT*) for conjugative delivery and a *pir*-dependent R6K origin. The R6K origin supports plasmid maintenance in specialized donor strains but prevents replication in recipient hosts lacking the π replication protein (Pir)^37^. We paired this with a conjugation-compatible Tn7 strategy^23^ for landing-pad integration. Once a compatible poly-*attB* landing pad is installed, the same recombinase and payload plasmids can be deployed across diverse bacteria without host-specific redesign.

We first use *Rhodobacter sphaeroides* to test whether conjugative delivery can overcome a defined barrier to SAGE. SAGE fails by electroporation in this host, a barrier not rescued by restriction endonuclease knockout (Fig. 1), but is resolved by cSAGE, which increases recovery of genomic integrants by at least five orders of magnitude relative to delivery of the same vectors by electroporation. We next show that cSAGE supports site-specific genomic integration across a diverse panel of bacteria, including several non-model PNSB strains, two *Pseudomonas* species, and *Rhodococcus jostii*. The toolkit also supports sequential multi-payload integration and marker recycling. Finally, we apply cSAGE to compare strain-specific differences in photosynthetic conversion of the lignin-derived aromatic *p*-coumarate to the polymer precursor *p*-vinylphenol across *Rhodopseudomonas palustris* strains, including the wastewater-derived environmental isolate P4. The same integrated pathway produced *p*-vinylphenol in P4 but not in the reference strain CGA009. Together, these results establish cSAGE as a portable platform for sequential genome engineering and comparative strain prototyping in genetically inaccessible bacteria.

**Figure 1.**
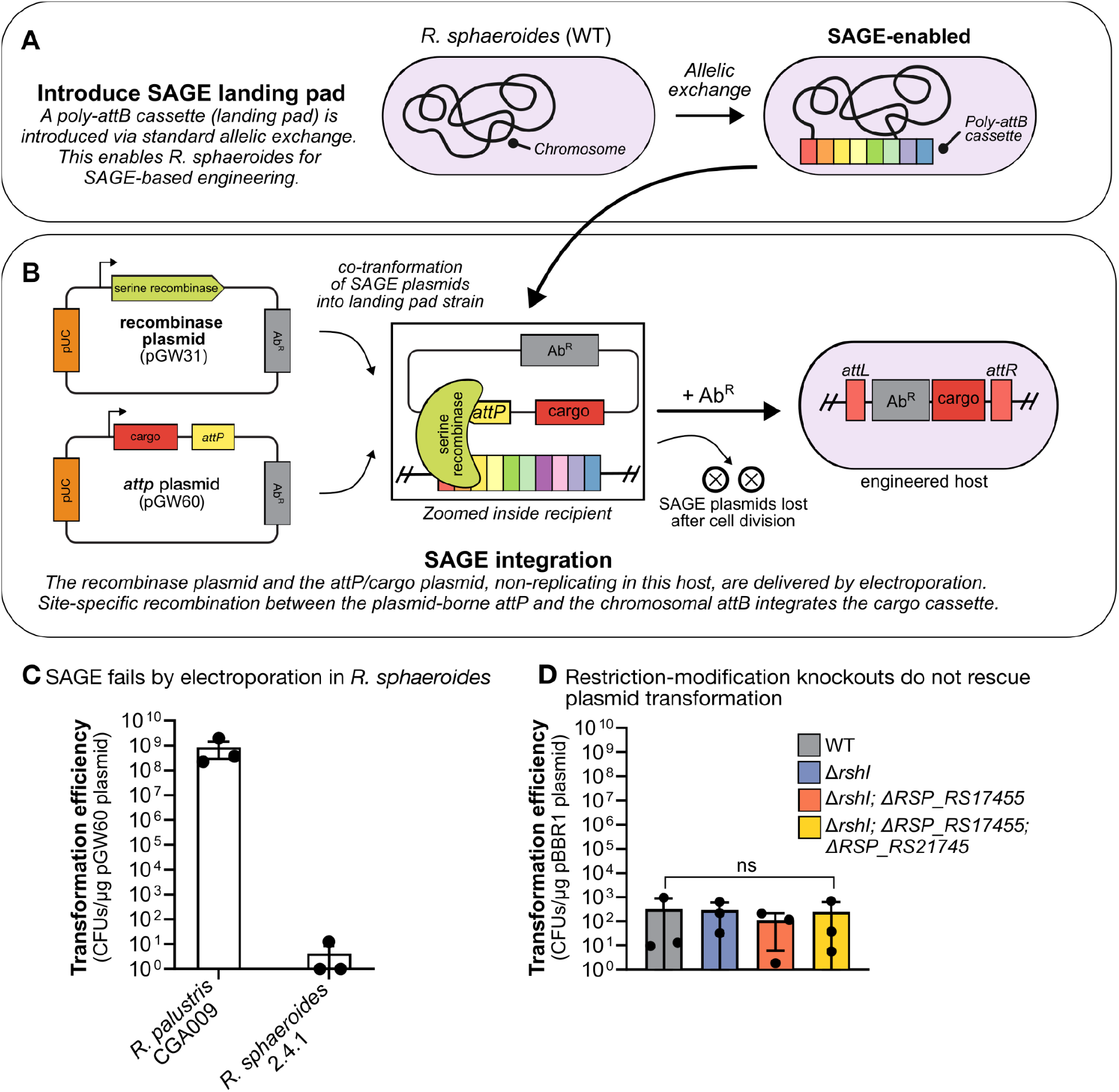
SAGE fails by electroporation in *Rhodobacter sphaeroides*. **(A)** Overview of landing pad installation. A poly-*attB* cassette containing multiple orthogonal *attB* sites is introduced into the *R. sphaeroides* chromosome at the *rshI* site by standard allelic exchange, generating a SAGE-enabled landing pad strain. **(B)** Overview of SAGE integration. The integrase plasmid (pGW31) carries a Bxb1 recombinase and pGW60 carries the cognate Bxb1 *attP* site. Both plasmids, non-replicating in this host, are co-transformed into the landing pad strain. The serine recombinase catalyzes site-specific recombination between the plasmid-borne *attP* and the chromosomal *attB*, integrating the cargo cassette and forming *attL* and *attR* junctions. The non-replicating plasmids are lost after cell division, leaving the engineered host with a single stably integrated cassette. **(C)** SAGE fails by electroporation in *R. sphaeroides*. Transformation efficiency (CFUs/µg pGW60 plasmid) following electroporation of *R. palustris* CGA009 (positive control) and *R. sphaeroides* 2.4.1 (*n* = 3; bars = mean ± s.d.). **(D)** Restriction-modification knockouts do not rescue electroporation efficiency. Transformation efficiency (CFUs/µg pBBR1 plasmid) in *R. sphaeroides* WT and restriction endonuclease knockout strains tested using the replicating plasmid pBBR1 (pCK524) (*n* = 3; bars = mean ± s.d.). *RSP_RS17455* and *RSP_RS21745* encode putative restriction endonucleases. The *P* values were determined by one-way ANOVA followed by a pairwise test with Bonferroni adjustment (ns, not significant).

## Results

### SAGE fails by electroporation in Rhodobacter sphaeroides, a barrier not rescued by restriction-modification knockout

Serine recombinase-assisted genome engineering (SAGE) enables single-copy, site-specific integration of genetic cargo through recombination between a chromosomally integrated poly-*attB* landing pad and a plasmid-borne *attP* site^26^. This system was previously validated in *Rhodopseudomonas palustris* CGA009 and several other bacterial species^26^. In the original workflow, an integrase expression plasmid and an *attP*/cargo plasmid are delivered by electroporation (Fig. 1A-B). *Rhodobacter sphaeroides* have a long history of use in bacterial photosynthesis research, yet remain resistant to standard transformation methods^7,38^. To extend SAGE to this host, we introduced a poly-*attB* landing pad cassette into the *R. sphaeroides* 2.4.1 genome by allelic exchange. Restriction-modification systems are a common barrier to foreign DNA uptake in transformation-resistant bacteria, and knockout of the *R. sphaeroides* type II restriction endonuclease RshI was previously reported to improve transformation efficiency^38,39^. We therefore integrated the landing pad cassette directly at the *rshI* locus. This approach simultaneously installed the poly-*attB* array and disrupted *rshI*, generating the Δ*rshI* landing-pad strain (Fig. 1A).

We then tested whether genetic payloads could be delivered to the landing pad strain using SAGE. We co-electroporated the integrase plasmid (pGW31) and *attP*/cargo plasmid (pGW60) into *R. palustris* CGA009 as a positive control and into the *R. sphaeroides* landing pad strain. We quantified transformation efficiency by selecting the cargo plasmid resistance marker (Fig. 1C). Electroporation yielded high transformation efficiency in *R. palustris* CGA009, consistent with prior reports^26^. In *R. sphaeroides*, however, transformants were recovered at or near the limit of detection, despite disruption of *rshI* in the landing pad strain. The most extensive published optimization of *R. sphaeroides* electroporation achieved an approximately tenfold improvement, reaching above 7,000 transformants per μg of DNA with a single replicating plasmid^39^. This improvement was unlikely to overcome the substantially greater barrier associated with delivering two non-replicating SAGE vectors. We therefore did not pursue further optimization of electroporation parameters for *R. sphaeroides* and instead investigated additional candidate restriction barriers.

To test whether additional, uncharacterized restriction-modification systems contributed to this barrier, we identified two further candidate type II restriction endonucleases encoded in the *R. sphaeroides* genome, RSP_RS17455 and RSP_RS21745. We generated a series of strains with sequential in-frame deletions (Supplementary Table 1) and compared their transformation efficiencies with that of wild-type *R. sphaeroides* by electroporation of pBBR1, a broad-host-range replicating plasmid^40^. We used pBBR1 rather than the non-replicating SAGE vectors because this single, autonomously replicating plasmid provided a higher baseline transformation efficiency and allowed us to evaluate the candidate restriction-endonucleases without the added complexity of delivering two non-replicating SAGE vectors (Fig. 1D).

Transformation efficiency did not improve with successive restriction-endonuclease knockouts. The knockout strains showed efficiencies comparable to that of wild-type *R. sphaeroides* (*P* = 0.86, one-way ANOVA; Fig. 1D). These results indicate that the three restriction endonucleases examined here are insufficient to explain the electroporation barrier in *R. sphaeroides*. Additionally, currently undefined mechanisms may therefore limit recovery following electroporation.

### Conjugation-based SAGE resolves the engineering bottleneck in Rhodobacter sphaeroides

Because disruption of the tested restriction endonucleases knockouts did not improve electroporation efficiency in *R. sphaeroides*, we evaluated conjugation as an alternative delivery route. Conjugation delivers DNA as a single-stranded intermediate directly through a mating pore^32–34^. This bypasses several steps required for electroporation, including membrane permeabilization and cytoplasmic entry of double-stranded DNA, that may limit uptake in this host^28^. Conjugation is also broadly compatible with Gram-negative bacteria and does not require host-specific preparation of electrocompetent cells, making it an attractive strategy for extending SAGE to transformation-resistant hosts^29–31^. Conjugative delivery has previously enabled genome engineering in other transformation-resistant bacteria facing similar barriers, including Tn7-based transposition^19,23^.

To enable conjugative delivery, we redesigned the SAGE plasmids by incorporating an RK2/RP4 *oriT*, generating the integrase plasmid pCK1205 and the *attP*/cargo plasmid pAR054 (Fig. 2A). We further restructured both plasmids into the Standard European Vector Architecture (SEVA) format, a modular design widely adopted for non-model bacterial hosts^35,36^ (Fig. 2A, Supplementary Fig. 1). We also replaced the replicon with R6K, a low-copy origin that requires the *pir* gene for replication^37^. The *pir*-dependent R6K origin lowers plasmid copy number and metabolic burden during cloning in *pir*⁺ *E. coli*^41^ and ensures the plasmids do not replicate in *R. sphaeroides* or other recipient hosts following conjugation^37^, so that only cells with a stable, integrated cargo cassette retain antibiotic resistance.

**Figure 2.**
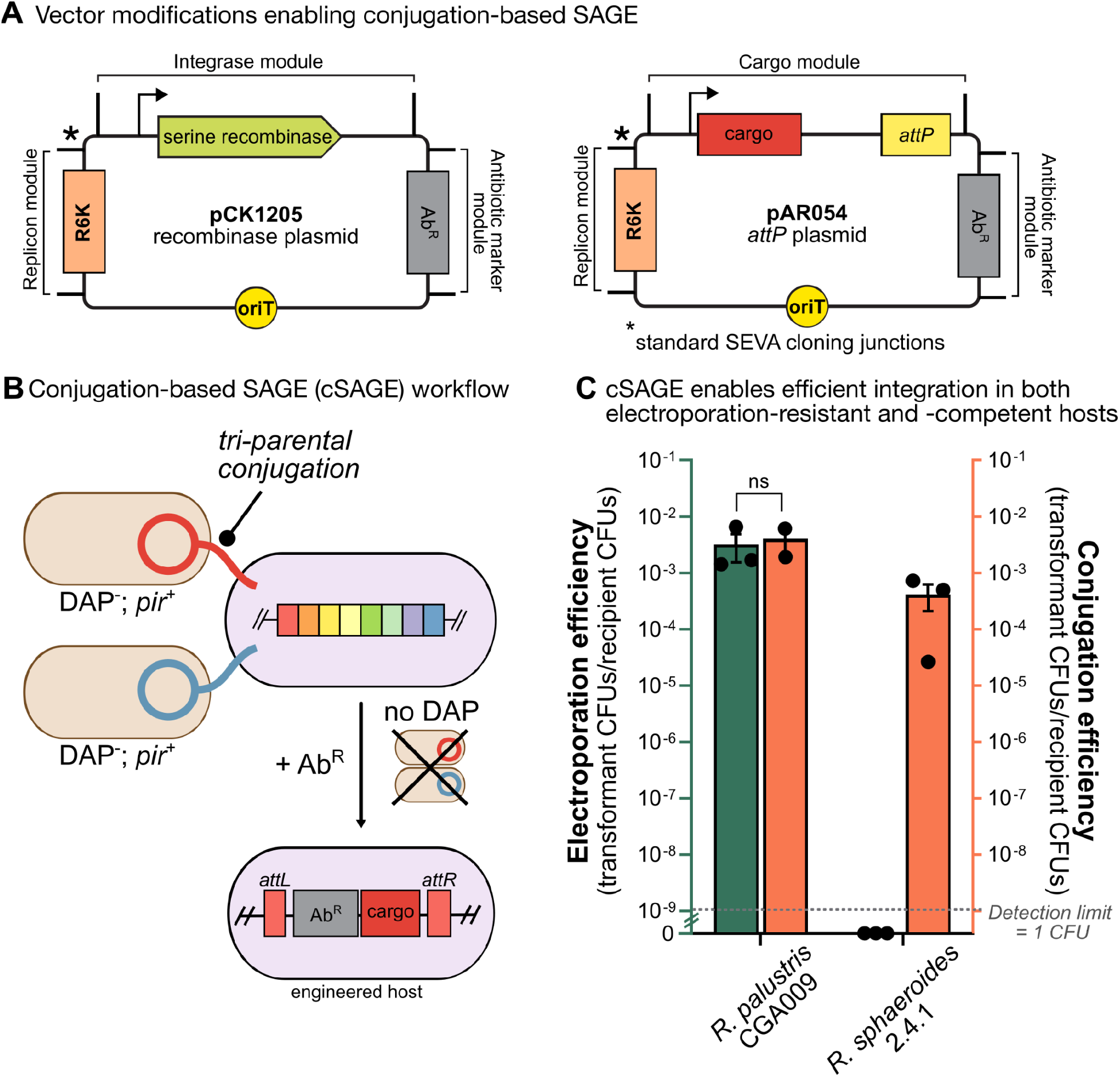
Conjugation-based SAGE (cSAGE) resolves the engineering bottleneck in *Rhodobacter sphaeroides*. **(A)** Vector modifications enabling cSAGE. SAGE plasmids were redesigned with a modular SEVA architecture, with each module flanked by standard SEVA cloning junctions (*). The integrase plasmid (left, pCK1205) and *attP*/cargo plasmid (right, pAR054) were made conjugation-compatible by addition of an RK2/RP4 origin of transfer (*oriT*). The replicon was changed to R6K, a low-copy origin that requires *pir* in trans for replication. **(B)** cSAGE workflow. Two DAP-auxotrophic *E. coli* donor strains, one carrying the recombinase plasmid and the other the *attP*/cargo plasmid, are mated with a SAGE-enabled recipient strain by tri-parental conjugation, transferring each plasmid via the mating pilus. Mating mixtures are then plated on medium lacking DAP and containing the antibiotic corresponding to the cargo plasmid’s resistance marker. This selects against donor growth, since the donors are DAP-auxotrophic, and for recipient cells with a stably integrated cargo cassette, since the non-replicating cSAGE plasmids are lost after division. **(C)** cSAGE enables efficient integration in both electroporation-resistant and -competent hosts. Electroporation efficiency (left, y-axis) and conjugation efficiency (right, y-axis) are shown for R6K-based cSAGE vectors in *R. palustris* CGA009 and *R. sphaeroides* 2.4.1. Conjugation efficiency and electroration efficiencies are expressed as transconjugant (transformant) CFUs per total recipient CFUs (*n* ≥ 2; bars = mean ± s.d.). Electroporation efficiencies are normalized to input DNA concentration. The *P* values were determined by Welch’s *t*-test (ns, not significant). Source data is provided as a Source Data File.

To deliver these plasmids, we used a tri-parental mating strategy (Fig. 2B). Two *E. coli* WM6026 donor strains, each *pir*⁺ and Δ*dapA* and therefore auxotrophic for diaminopimelic acid (DAP)^42^, were used: one carrying the integrase plasmid and the other carrying the *attP*/cargo plasmid. Both donor strains were co-cultured with the SAGE-enabled *R. sphaeroides* recipient, allowing conjugative transfer of both plasmids into the same recipient cell via the mating pilus. Mating mixtures were then plated on medium lacking DAP and containing the antibiotic corresponding to the cargo plasmid’s resistance marker. This selects against both donor strains, which cannot grow without DAP, and for recipient cells with stable, site-specific integration of the cargo cassette, since the non-replicating SAGE plasmids are otherwise lost after cell division.

Using this conjugation-based approach (cSAGE), we compared electroporation and conjugation efficiency (both of which were normalized to total recipient CFUs) directly in *R. palustris* CGA009 and *R. sphaeroides*, using the same R6K-based cSAGE vectors for both delivery methods to isolate delivery method as the sole variable (Fig. 2C). Conjugation and electroporation gave comparable recovery frequencies in *R. palustris* CGA009 (*P* = 0.71, Welch’s *t*-test; Fig. 2C) but conjugation yielded at least five orders of magnitude more recovered integrants than electroporation in *R. sphaeroides* (Fig. 2C). In *R. sphaeroides*, cSAGE efficiently and reproducibly achieved transconjugant recovery at frequencies of ∼10^−4^ per recipient cell (1 in 10,000), whereas electroporation efficiency fell below our limit of detection (≤10^−9^). Under the matched tri-parental mating conditions used here, the tested mini-Tn7 configuration yielded no recoverable transconjugants in *R. sphaeroides* (Supplementary Fig. 2). Published Tn7 workflows using an equivalent tri-parental mating scheme have similarly failed to recover transconjugants^19^. In the published study, reliable recovery required either an additional conjugal helper strain or a recipient engineered to overexpress the Tn7 transposase, with mean frequencies of 10^−4^ to 10^−3^ across the tested configurations^19^. cSAGE achieves comparable transconjugant recovery (∼10^−4^ per recipient cell) using an unmodified, standardized tri-parental mating scheme, without the additional strain engineering that the best available Tn7 benchmark required. This comparison indicates that the advance is attributable specifically to the cSAGE vector and delivery design, rather than to conjugation-based delivery in general.

### Conjugation-based SAGE enables efficient genomic integration across diverse bacteria

To evaluate the portability of cSAGE beyond *R. sphaeroides*, we assembled a diverse panel of PNSB from the Alphaproteobacteria and Betaproteobacteria (Table 1)^43,44^. For *R. palustris* P4, *Rhodospirillum rubrum*, and *Rubrivivax gelatinosus*, we integrated the poly-*attB* landing pad using a transposon-based strategy^23^ (Fig. 3A). Tn7 systems have been widely adopted for single-copy gene integration in non-model organisms, but transposition immunity restricts integration to a single event per *attTn7* site^23^, precluding iterative use of Tn7 itself for additional integrations^45^. By using Tn7 only once to install a poly-*attB* landing pad containing ten orthogonal *attB* sites, we can enable iterative engineering at this locus. We introduced a single standardized delivery plasmid (pRC414) to SAGE-enable multiple hosts without redesigning integration vectors for each species. pRC414 itself carries an RK2/RP4 *oriT* (Fig. 3A) and was delivered to each recipient by conjugation, consistent with our conjugation-based delivery strategy (Fig. 2B).

**Figure 3.**
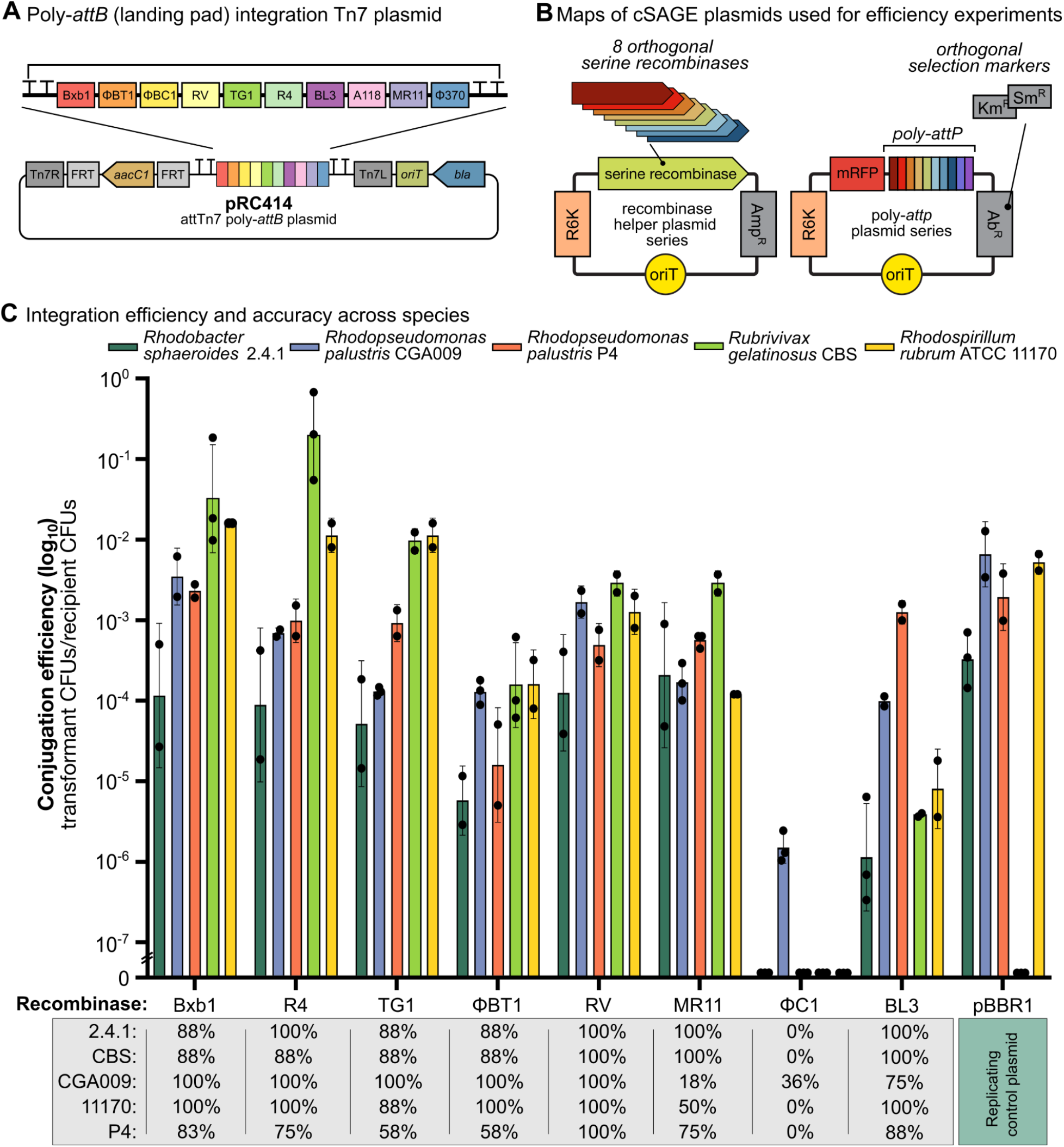
cSAGE recombinase benchmarking across diverse hosts. **(A)** Design of the Tn7-based poly-*attB* landing pad delivery plasmid. The genomic landing pad consists of ten orthogonal serine recombinase attachment sites (*attB*), flanked by transcriptional terminators to prevent transcriptional readthrough and interference with neighboring loci. A Tn7 delivery vector (pRC414) was constructed to integrate the poly-*attB* cassette into Gram-negative bacterial genomes. The *aacC1* gentamicin resistance marker is flanked by FRT sites for optional removal via Flp recombinase. **(B)** Maps of recombinase and poly-*attP* plasmids used for recombinase screening. Integration was tested by tri-parental conjugation, delivering two non-replicating plasmids: (i) a poly-*attP* cargo plasmid carrying a constitutively expressed mRFP reporter and one of eight orthogonal *attP* sites, and (ii) an integrase expression plasmid encoding the cognate serine recombinase (Bxb1, ϕBT1, ϕC1, RV, TG1, R4, BL3, or MR11). Two additional landing pad sites, A118 and ϕ370, are included for future expansion but were not tested in this screen. **(C)** Recombinase activity and fidelity across purple nonsulfur bacteria (PNSB). Conjugation efficiencies (number of transconjugant colonies per recipient CFU; log₁₀ scale) are shown for eight recombinases tested in *Rhodobacter sphaeroides* 2.4.1, *Rhodopseudomonas palustris* CGA009, *R. palustris* P4, *Rubrivivax gelatinosus* CBS, and *Rhodospirillum rubrum* ATCC 11170. A replicating control plasmid (pBBR1) was used to benchmark conjugation efficiency (*n* ≥ 2; bars = mean ± s.d.). The table below summarizes integration accuracy, defined as the proportion of colonies with correct on-target integration confirmed by colony PCR (*n* ≥ 8; bars = mean ± s.d.). Source data is provided as a Source Data File.

**Table 1.** Bacteria used in this study.

| Organism | Classification | Origin | Primary bioproduction use cases |
| --- | --- | --- | --- |
| *Rhodopseudomonas palustris* CGA009 | Alphaproteobacteria | Soil | Aromatic compound valorization<sup>54–56,82</sup> |
| *Rhodopseudomonas palustris* P4 | Alphaproteobacteria | Anaerobic digester | Fermentative CO oxidation and hydrogen production<sup>43,60,61</sup> |
| *Rhodobacter sphaeroides* 2.4.1 | Alphaproteobacteria | Freshwater | Isoprenoid biosynthesis and biohydrogen production<sup>6,7,13</sup> |
| *Rhodospirillum rubrum* ATCC 11170 | Alphaproteobacteria | Freshwater | Anaerobic syngas fermentation<sup>9</sup> |
| *Rubrivivax gelatinosus* CBS | Betaproteobacteria | Soil | Photosynthetic fermentation of syngas and organic acids<sup>83</sup> |
| *Rhodococcus jostii* RHA1 | Actinomycetes | Soil | Lignin depolymerization and conversion of aromatic compounds<sup>49,75</sup> |
| *Pseudomonas sorgoleonovorans* SO81 | Gammaproteobacteria | Rhizosphere | Plant-growth promotion and utilization of plant-derived carbon sources<sup>48</sup> |
| *Pseudomonas putida* KT2440 | Gammaproteobacteria | Soil | Metabolic engineering platform for aromatic compound valorization<sup>46,47</sup> |

Using this panel of five PNSB strains, we screened eight orthogonal serine recombinases: Bxb1, R4, TG1, ϕBT1, RV, MR11, ϕC1, and BL3. Each recombinase was delivered by tri-parental conjugation as in Fig. 2B. cSAGE produced detectable transconjugants in all five PNSB strains, and for nearly all recombinase-host combinations (Fig. 3C). To enable this comparison across recombinases and hosts within a consistent genetic context, we employed a poly-*attP* reporter plasmid series, each carrying one of eight *attP* sites paired with a constitutively expressed mRFP reporter (Fig. 3B)^26^. When paired with the cognate recombinase expression plasmid, integration is directed specifically to the matching *attB* site, resulting in chromosomal insertion of the mRFP cassette, confirmed by colony PCR. As a benchmark, we also measured conjugation efficiency using a pBBR1 control plasmid.

Both integration efficiency and accuracy influence the practical utility of a recombinase. We quantified conjugation efficiency as the frequency of transconjugant colonies per viable recipient cell and integration accuracy as the fraction of screened colonies with on-target integration. Bxb1, R4, and TG1 consistently yielded high conjugation efficiency across all strains, with some values comparable to pBBR1 control (Fig. 3C). The remaining ϕC1, ϕBT1, RV, and BL3 recombinases showed variable conjugation efficiency across species, with ϕC1 yielding detectable transconjugants only in *R. palustris* CGA009. Bxb1, TG1, R4, and RV were also among the most accurate recombinases tested across all five species. The remaining recombinases showed more variable integration accuracy compared to Bxb1, TG1, R4, and RV.

No transconjugants were recovered in *R. gelatinosus* CBS using the replicating pBBR1 control, whereas cSAGE integrations were readily obtained in this host (Fig. 3C). To assess whether this portability extends beyond PNSB, we evaluated cSAGE in phylogenetically distinct bacteria, including *Pseudomonas putida* KT2440^46,47^, *Pseudomonas sorgoleonovorans* SO81^48^, and *Rhodococcus jostii* RHA1^49^, using the three highest-performing recombinases identified in PNSB (Bxb1, R4, and TG1) (Supplementary Fig. 3). Electroporation of *P. sorgoleonovorans* was previously found to be highly inefficient, requiring conjugation for gene deletion and DNA integration^48^. We observed detectable integration across all three hosts although transconjugant recovery frequency and integration accuracy varied among recombinase-host combinations. While this dataset is not intended as a comprehensive host-range analysis, cSAGE supported genomic integration across four bacterial classes spanning two phyla.

### Iterative genome engineering workflows enabled by cSAGE

Iterative strain engineering in non-model bacteria involves repeated rounds of genomic integration to incrementally assemble complex or burdensome genetic programs, such as multi-gene metabolic pathways that cannot be introduced in a single step^15^. In practice, such workflows are limited by the repertoire of effective selectable markers^21,50^ due to intrinsic resistance to antibiotics^13^, cross-resistance conferred by some antimicrobial resistance genes ^51^, and inconsistent performance of counterselection systems across species^50^. As a result, many existing approaches combine sequential rounds of integration with intermediate marker or backbone removal to enable marker reuse, adding experimental steps and imposing additional host-specific requirements^22^. These limitations are particularly prevalent in non-model organisms such as PNSB, where counterselection systems are often unavailable, unreliable, or require extensive characterization alongside selection markers^52^. Engineering workflows that enable rapid construction of multi-payload strains without mandatory marker recycling, while retaining optional marker excision, remain scarce. To address this need, we developed two conjugation-compatible workflows: a multi-payload integration toolkit that enables sequential construction of marked strains without counterselection, and a ϕC31-based marker-recycling strategy that supports removal of antibiotic resistance markers where compatible counterselection systems exist.

We tested whether multiple payloads could be integrated sequentially into cognate sites within the poly-*attB* landing pad without cross-recombination or interference. We designed payload plasmids containing distinct *attP* sites matched to orthogonal *attB* loci to enable independent recombinase-mediated integration events. We constructed three orthogonal serine recombinase systems–Bxb1, R4, and TG1–based on their strong performance and paired each with a unique antibiotic-resistance marker (Km^R^, Sm^R^, or Gm^R^) for independent selection (Fig. 4A). As a proof of concept, we introduced orthogonal fluorescent reporters (mRFP, sfGFP, and mTagBFP2) as cargos into each of the three attachment sites. We performed sequential rounds of integration in both *R. palustris* CGA009 and *R. sphaeroides*, selecting only for the newly introduced antibiotic marker at each step and without intermediate excision of previously integrated backbones. Despite the introduction of repetitive R6K backbone sequences from prior integrations, fluorescence from all three reporters was retained through the successive rounds of integration (Fig. 4B-C; Supplementary Fig. 5).

**Figure 4.**
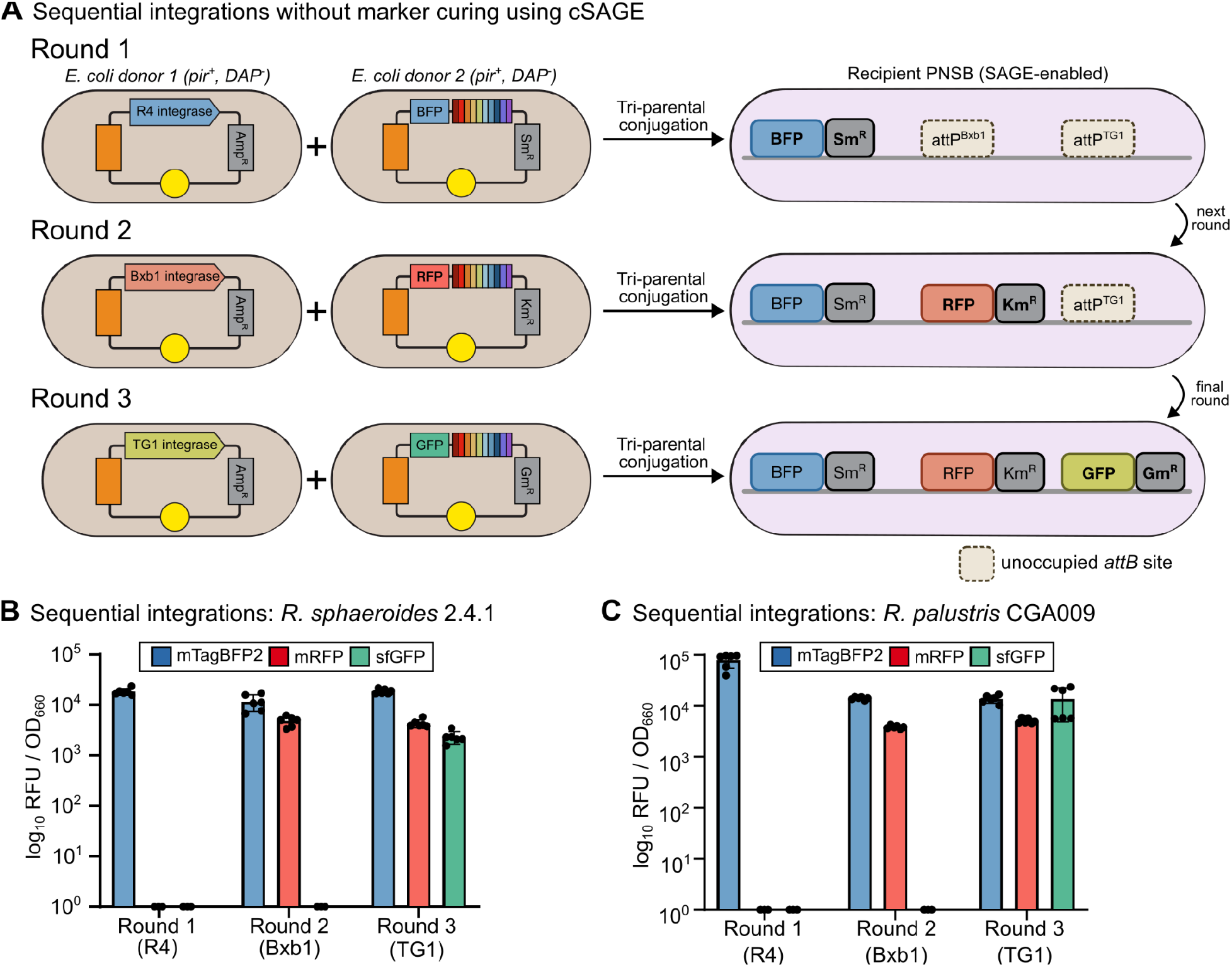
Sequential multi-payload integration via cSAGE without marker curing. (**A**) Three rounds of tri-parental conjugation deliver orthogonal fluorescent reporter cassettes sequentially into a SAGE-enabled PNSB recipient. In each round, two *E. coli* donor strains (pir⁺, DAP^−^) carry a non-replicating integrase expression plasmid and a cognate non-replicating *attP*/cargo plasmid, respectively. Round 1 delivers R4 integrase and a BFP cargo with a spectinomycin resistance marker (Sm^R^); Round 2 delivers Bxb1 integrase and an RFP cargo with kanamycin resistance (Km^R^); Round 3 delivers TG1 integrase and a GFP cargo with gentamicin resistance (Gm^R^). Each round selects only for the newly introduced antibiotic marker, with no selection for or excision of previously integrated cassettes. Unoccupied *attB* sites remaining in the landing pad after each round are shown as dashed boxes. (**B-C**) Fluorescence quantification after each integration round in *Rhodobacter sphaeroides* 2.4.1 (**B**) and *Rhodopseudomonas palustris* CGA009 (**C**). Fluorescence from all reporters present in the strain (newly integrated and previously integrated) was quantified at each round to assess expression and confirm absence of cassette dropout (*n* = 6; bars = mean ± s.d.).

While the multi-payload integration strategy enables strain construction without backbone excision, the generation of antibiotic-resistance-free strains remains important for many downstream applications. In the SAGE system, ϕC31 integrase catalyzes excision of plasmid backbones flanked by *attP* and *attB* sites following payload integration, leaving behind only the desired cargo and a small recombination scar (*attL*) with minimal expected impact on host physiology^26^. However, delivery of ϕC31 integrase in the original implementation relied on either a replicative mSF^ts1^ plasmid (a *Pseudomonas*-specific origin^53^) or a non-conjugative pUC-derived plasmid^26^. These delivery systems have limited applicability in PNSB where mSF^ts1^ does not replicate and electroporation is inefficient. To evaluate marker recycling and backbone excision in a fully conjugation-based workflow, we reimplemented the ϕC31 integrase helper plasmid in the conjugative R6K vector. We constructed a poly-*attP* target plasmid containing constitutively expressed *mRFP* and a *sacB* gene for counterselection (Fig. 5A). We integrated this *sacB*-containing cargo into *R. palustris* CGA009 and *R. sphaeroides* using a Bxb1-based cSAGE protocol. Strains with the chromosomally integrated cargo then underwent diparental conjugation to deliver the ϕC31-encoding R6K plasmid. Following conjugation and sucrose counterselection, we observed marker curing in both species (Fig. 5B-C). In *R. palustris* CGA009, 16 of 16 colonies yielded the expected ∼1.5 kb PCR product, consistent with precise excision of the plasmid backbone (Fig. 5B). Similarly, in *R. sphaeroides*, 14 of 16 colonies yielded the expected PCR product; no diagnostic amplicon was obtained from the remaining two colonies, leaving their curing status unresolved.

**Figure 5.**
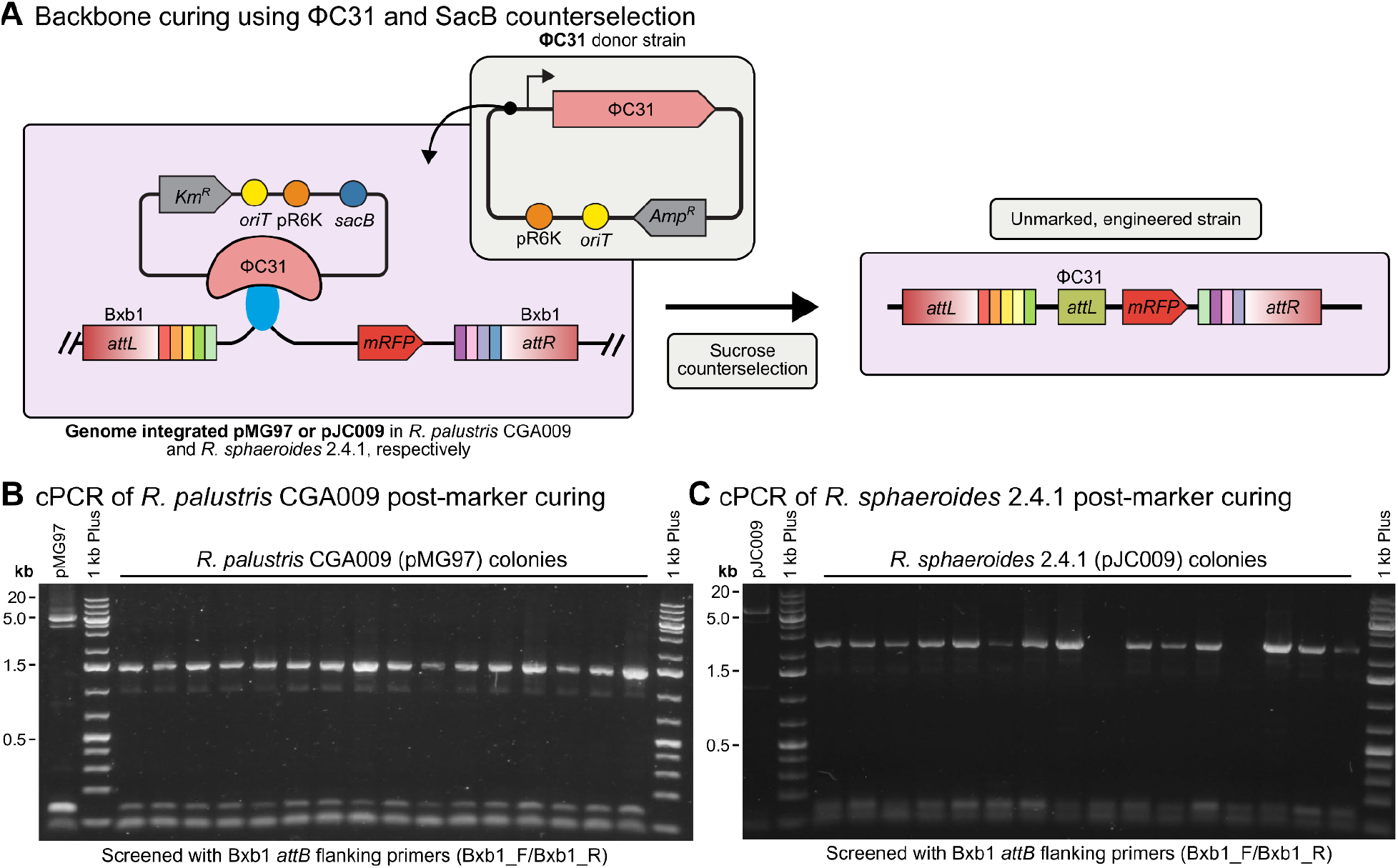
Marker recycling in cSAGE using ϕC31 integrase-mediated excision and sucrose counterselection. (**A**) ϕC31-based marker curing strategy. Genome-integrated poly-*attP* plasmids (pMG097 or pJC009) containing an *mRFP* reporter and a *sacB* counterselection cassette were flanked by ϕC31 *att* sites and delivered to *Rhodopseudomonas palustris* CGA009 or *Rhodobacter sphaeroides* 2.4.1. A non-replicating ϕC31 integrase expression plasmid (pSG010) was subsequently introduced by conjugation to catalyze site-specific recombination between the ϕC31 *att* sites, excising the plasmid backbone and antibiotic resistance marker. The resulting markerless strain retains only the integrated cargo and minimal *attL*/*attR* scars. Following ϕC31-mediated recombination, sucrose counterselection was applied to remove residual cells retaining the *sacB* cassette/backbone. (**B-C**) Verification of marker excision. Colony PCR using primers flanking the Bxb1 *attB* integration site were used to distinguish between uncured (larger amplicon) and cured (smaller amplicon) strains. Expected band sizes: pMG097, 5125 bp (uncured) and 1461 bp (cured); pJC009, 6400 bp (uncured) and 2313 bp (cured). Thermo Scientific GeneRuler 1KB Plus DNA Ladder was used as a reference.

The multi-payload integration toolkit supports reliable, counterselection-free construction of strains carrying multiple integrated payloads, making it particularly well suited for workflows where rapid pathway assembly, testing, and comparison are prioritized. In parallel, ϕC31-mediated marker recycling provides an efficient route to generate antibiotic-resistance-free strains when compatible counterselection systems are available. These capabilities establish a flexible engineering framework for strain construction that supports both rapid prototyping and marker-free strain construction.

### Light-powered aromatic bioproduction in purple nonsulfur bacteria

PNSB represent one of the most metabolically versatile groups of microorganisms known, capable of thriving under anaerobic conditions while using infrared light energy and a broad range of inorganic and organic compounds as energy and carbon sources^5^. *R. palustris*, in particular, can oxidize diverse substrates, including CO_2_, H_2_, and organic acids, making it a promising chassis for circular bioeconomy applications^6^. The ability of *R. palustris* to catabolize lignin-derived aromatics under anoxygenic conditions^54–56^ further distinguishes this species as a candidate for valorizing industrial waste streams such as lignocellulosic hydrolysates^10,56^ and wastewater effluents^8^. However, this metabolic versatility is underpinned by complex regulatory networks that respond to light availability, O_2_ tension, and nutrient status to control distinct carbon assimilation and redox balancing pathways^57^. Effective bioproduction in PNSB, therefore, requires not only understanding their metabolic potential but also genome engineering tools that enable predictable control of heterologous pathway expression across species.

For well-studied PNSB such as *R. palustris* CGA009, genome-scale metabolic models (GEMs) provide a useful framework for predicting metabolic fluxes and evaluating production potential^58^. However, GEM construction is labor-intensive, requires extensive curation and experimental validation, and models are often not readily transferrable even between closely related strains^59^. This limitation becomes particularly apparent when working with environmental isolates. For example, we recently sequenced and annotated the genome of the wastewater isolate *R. palustris* P4 (GenBank accession number CM151374.1). P4 has a previously characterized ability to couple carbon monoxide oxidation to hydrogen production through the water-gas shift reaction^43,60,61^, but its capacity to degrade aromatic compounds had not been previously characterized. To our knowledge, P4 had also not previously been genetically engineered, making this study the first demonstration of genome engineering in this strain. Comparative genomic analysis revealed numerous strain-level differences relative to CGA009, complicating *a priori* prediction of how native metabolism would interact with an engineered pathway (Supplementary Table 6 and 7). In this context, cSAGE provides an empirical approach to evaluate metabolically distinct but poorly characterized environmental isolates for bioproduction. Rather than optimizing a pathway within a single host, our goal was to deploy the same genetic constructs across multiple hosts to identify productive strain backgrounds.

To illustrate this capability, we focused on *R. palustris* strains as hosts for aromatic bioconversion. *R. palustris* possesses well-characterized pathways for uptake and catabolism of *p*-coumarate (*p*-CA) and related aromatics, including a conserved coumarate degradation operon (the *CouAB* genes and associated transport systems^54,62^). Because substrate uptake and native catabolic capacity are prerequisites for conversion, *R. palustris* provides a relevant chassis for evaluating aromatic bioproduction. We therefore integrated a codon-optimized *BaPAD* expression cassette encoding a phenolic acid decarboxylase from *Bacillus atrophaeus*. BaPAD converts *p*-CA to *p*-vinylphenol (*p*-VP)^63,64^, a precursor for polymers^65^, adhesives^66^, and photolithography materials^67^ (Fig. 6A). Because *R. palustris* strains differ in their genomic content (Supplementary Tables 6 and 7), we hypothesized that host-specific physiology could strongly influence bioproduction outcomes even when the heterologous pathway is introduced.

**Figure 6.**
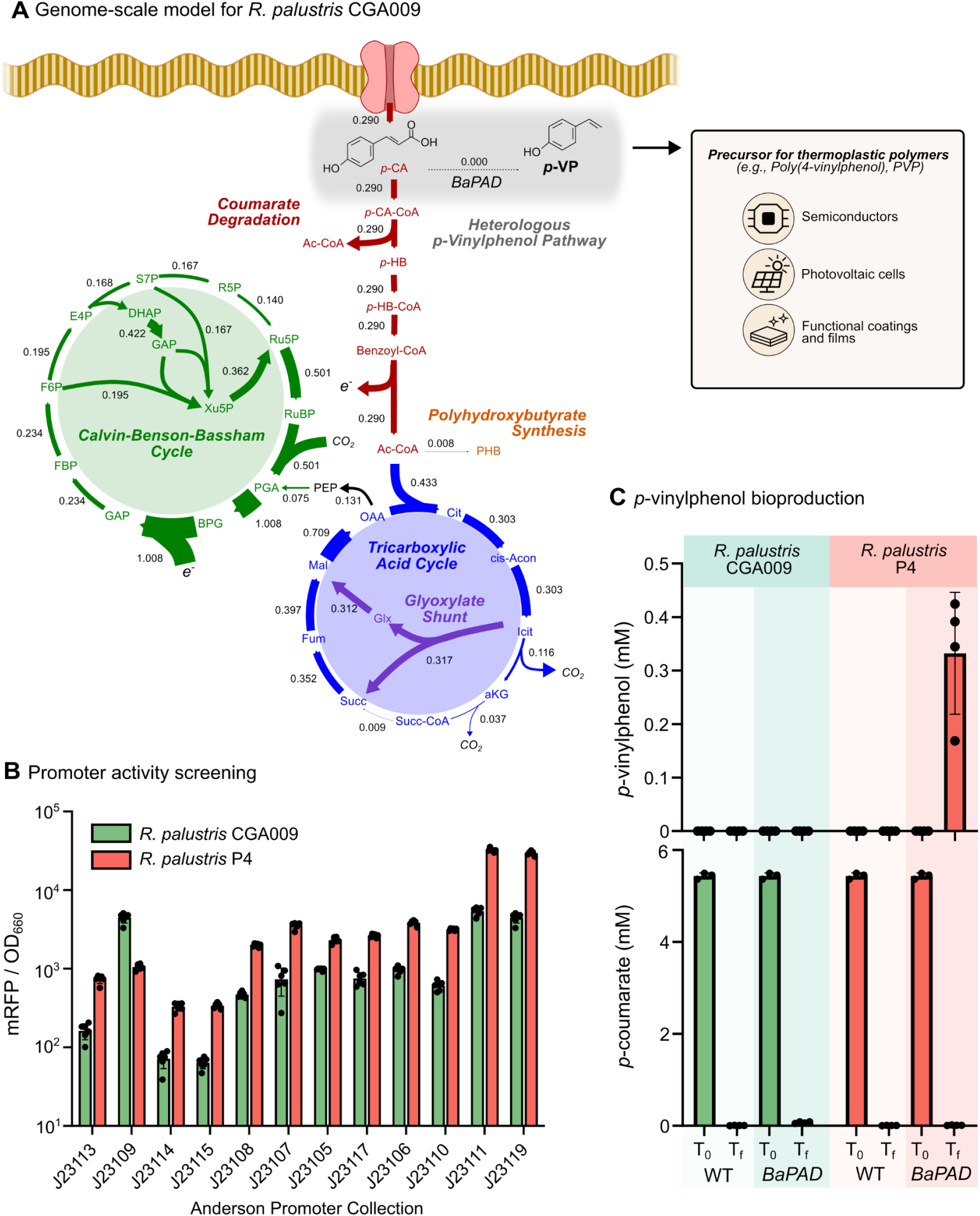
Photosynthetic bioproduction of *p*-vinylphenol in engineered PNSB. (**A**) Simulations of an iAN1128-based genome-scale model of engineered phenolic acid decarboxylase (PAD)-expressing *R. palustris* CGA009 grown photoheterotrophically on *p*-coumarate (*p*-CA). (**B**) Promoter activity across hosts. Fluorescence of integrated promoter constructs in *R. palustris* P4 and CGA009. Data show relative fluorescence units (RFU) normalized to optical density (OD₆₆₀) for replicate colonies from transformation plates (*n* = 6; bars = mean ± s.d.). (**C**) *p*-vinylphenol production and *p*-CA consumption in *R. palustris* P4 and CGA009. Metabolite concentrations (*p*-VP and *p*-CA) were measured at inoculation (T₀) and after cultivation (T_f_) for both wild-type (WT) and engineered (BaPAD) strains. *p*-VP is normalized to biomass at the time of sampling (*n* ≥ 3; bars = mean ± s.d.). Source data is provided as a Source Data File.

Before integrating BaPAD, we used cSAGE to benchmark promoter activity across the two *R. palustris* strains using a set of synthetic constitutive promoters characterized in *E. coli* and other Gram-negative bacteria^68^ (Fig. 6B; Supplementary Fig. 6). Promoter characterization is important for controlling heterologous gene expression. Although the rank correlation of these synthetic promoters is generally conserved in previously characterized species^68^, their activity can vary among hosts and is best determined experimentally. Using mRFP as a reporter, we identified BBa_J23110 as a medium-strength promoter within each hosts’ promoter panel, though its absolute expression level differed between hosts. We then used cSAGE to integrate the BBa_J23110::BaPAD expression cassette. The engineered strains were cultivated photoheterotrophically with *p*-CA as the sole carbon source (Fig. 6C, Supplementary Fig. 7).

Despite complete consumption of *p*-CA substrate, the CGA009_BaPAD strain produced no detectable extracellular *p*-VP (Fig. 6C, Supplementary Fig. 7). This result is consistent with native *p*-CA metabolism limiting net product accumulation. Flux simulations using the iAN1128 GEM adapted for CGA009_BaPAD^58^ predicted no net flux toward *p*-VP production and instead routed carbon through native aromatic degradation pathways and into central metabolic intermediates such as acetyl-CoA. In contrast, *R. palustris* P4_BaPAD produced measurable amounts of *p*-VP, converting approximately 6% of the supplied *p*-CA to product (Fig. 6C). The *p*-CA catabolic operon has an identical nucleotide sequence in CGA009 and P4 (Supplementary Fig. 8). Comparative analysis of KEGG orthologies identified additional functions in P4 associated with phenylpropanoid metabolism, redox processes, and transport. Differences in redox balance, pathway regulation, product turnover or export could therefore contribute to the observed divergence in *p*-vinylphenol accumulation between strains (Supplementary Tables 6 and 7).

Together, these results demonstrate that host physiology can strongly influence pathway performance, even among closely related strains. More broadly, this experiment illustrates how cSAGE enables direct evaluation of bioproduction pathways across hosts using empirical performance data. By enabling stable genome integration across hosts, cSAGE reduces confounding effects associated with plasmid copy number and genetic instability^13^. This capability provides a practical route to identify productive strain backgrounds early in the strain development pipeline, particularly for non-model organisms for which predictive models are unavailable or insufficient.

## Discussion

Poor transformation efficiency remains one of the most common barriers to genome engineering in microbial synthetic biology^14^. Conventional domestication strategies scale poorly because each new host typically requires constructing bespoke allelic exchange vectors, identifying compatible origins of replication, and validating genetic parts from scratch^15,22^. This challenge is not limited to poorly characterized organisms. In *R. sphaeroides*, targeted disruption of candidate restriction endonucleases failed to improve electroporation efficiency (Fig. 1). Conjugative delivery of the same vectors instead increased recovery of genomic integrants by at least five orders of magnitude (Fig. 2), indicating that the delivery route was a major limitation. Benchmarking against a standard Tn7-based conjugation workflow^19^ under matched conditions further showed that cSAGE achieved comparable or better transconjugant recovery (Fig. 2) without the additional donor or recipient engineering the published protocol required, indicating that the improvement reflects the cSAGE vector and delivery design specifically, rather than conjugation-based delivery in general. cSAGE couples conjugative DNA delivery with serine recombinase-mediated genomic integration using a *pir*-dependent R6K vector backbone. Once a compatible landing pad is installed, the same pre-built recombinase and payload plasmids can be used in new organisms with minimal vector development. This includes previously uncharacterized environmental isolates such as *R. palustris* P4, for which no genetic tools had been reported prior to this work. cSAGE relies on Tn7 transposition or allelic exchange for the one-time installation of its landing pad. These complementary methods remain effective for this initial step but require host- and locus-specific optimization (e.g., homology arm design) for each new target (Table 3).

The serine recombinases varied in performance across hosts (Fig. 3), despite being expressed from identical promoters and delivered using the same conjugation workflow. This variation suggests that performance is influenced by intrinsic sequence preferences and host genomic context^69,70^. RV achieved 100% on-target integration across all five PNSB despite low overall transconjugant recovery frequencies (Fig. 3), indicating that integration accuracy and recovery frequency are separable performance measures. Applying motif-scanning or pseudo-*attB* site prediction^71^ approaches to bacterial genomes could help explain these differences and guide recombinase selection, although empirical validation will remain important when porting cSAGE to new hosts.

Among the PNSB tested, *R. palustris* CGA009 and P4 were the most closely related, yet they exhibited divergent *p*-VP production and accumulation phenotypes (Fig. 6C). Although both CGA009 and P4 possess the *p*-CA catabolic operon (Supplementary Fig. 8), P4 produced measurable *p*-VP, whereas no extracellular p-VP was detected in CGA009. Reporter output from J23110 constructs integrated at the respective landing pads was higher in P4 than in CGA009 (*P* < 0.0001, one-way ANOVA; Fig. 6B), raising the possibility that differences in promoter activity and BaPAD enzyme expression levels contributed to the difference in *p*-VP titer. However, the approximately fivefold difference in reporter expression is unlikely to fully explain the contrasting production phenotypes. CGA009 produced no *p*-VP despite complete consumption of *p*-CA (Fig. 6C). This result is consistent with GEM simulations predicting zero net flux toward *p*-VP in CGA009 regardless of BaPAD expression level, with competing native catabolic pathways consuming the available flux upstream of the heterologous step (Fig. 6A). The underlying basis for this divergence remains unresolved, but comparative genomic analysis identified additional functions in P4 compared to CGA009 that are related to phenylpropanoid metabolism, redox processes, and transport (Supplementary Tables 6 and 7). These differences could contribute to the observed variation in net *p*-VP production.

Several non-mutually exclusive mechanisms could account for the distinct production phenotype. Differences in *p*-CA import or CoA ligation kinetics could alter the balance between BaPAD-mediated decarboxylation and native catabolic flux. CGA009 encodes two redundant, high-affinity *p*-CA transport systems^72^, and comparative genomic analysis indicates that P4 differs in transport-related loci (Supplementary Tables 6 and 7). More rapid uptake and CouB-mediated CoA ligation in CGA009 could reduce the pool of free *p*-CA available to BaPAD^62^. Differences in regulatory control of the coumarate degradation (*cou*) operon could similarly affect how quickly native catabolic flux is activated. In CGA009, this operon is controlled by the MarR-family repressor CouR and derepressed specifically by *p*-coumaroyl-CoA^62^. Strain differences in *p*-VP turnover, tolerance, or export may also contribute to the difference in extracellular product accumulation. Vinylphenol derivatives can be more toxic to Gram-negative bacteria than their parent phenolics^73^, making product tolerance and export relevant targets for further investigation. Distinguishing among these possibilities will require targeted experiments, including measurements of intracellular intermediates and pathway flux, together with transporter knockouts and *in vitro* enzyme assays^64^. Directly deleting competing catabolic genes could, in principle, redirect flux toward *p*-VP, but also narrow the aromatic substrate range that makes PNSB attractive as bioproduction hosts. The naturally genome-reduced *Rhodopseudomonas* strain DSM127 has substantially fewer genes associated with aromatic-compound biodegradation, illustrating how genome reduction can narrow selected metabolic capabilities^74^. Targeted, additive engineering approaches such as cSAGE, which leave native metabolism intact, may therefore provide a useful strategy for metabolically versatile organisms such as PNSB, where the functional consequences of pathway deletion are not always predictable.

By combining portable genome engineering with photosynthetic chassis, cSAGE provides genetic access to organisms capable of valorizing underutilized carbon streams. The ability of PNSB to integrate CO₂ fixation, aromatic catabolism, and phototrophic growth positions them as promising platforms for sustainable bioproduction from lignin-derived aromatics, organic acids, and syngas^6,7^. The *p*-VP route demonstrated here illustrates this potential. BaPAD-catalyzed decarboxylation removes only the carboxyl group from *p*-CA, providing a direct route from a lignin-derived aromatic to a value-added product^56,63^. The multi-payload integration and promoter-characterization workflows developed here lay the groundwork for systematic strain engineering and integration with metabolic and systems-level models^11,58^. Beyond PNSB, cSAGE supported genomic integration in the Gram-positive *R. jostii* RHA1 and the Gram-negative bacteria *P. putida* KT2440 and *P. sorgoleonovorans* SO81 (Supplementary Fig. 3), demonstrating portability across phylogenetically distinct hosts relevant to bioremediation^75^, bioproduction^47^, and agriculture^48^. The standardized integration architecture could also complement CRISPR-based regulation in design-build-test-learn cycles by providing stable installation of pathway and control modules^76,77^, and facilitate synthetic pathway construction^78^, and genetic code expansion^79^. Applying the same genetic constructs across strain backgrounds could generate controlled, strain-resolved data for refining mechanistic and machine-learning models, particularly when limited chassis-specific knowledge and training data constrain prediction in non-model bacteria. By enabling common genetic designs to be installed across candidate hosts, cSAGE provides a practical route to expand genetic access and identify productive strain backgrounds among non-model bacteria.

## Methods

### Plasmid and strain construction

Plasmids used in this study are listed in Supplementary Table 2. Key plasmids used in this study, including Addgene IDs, are listed in Table 2. All cSAGE plasmids were assembled in the Standard European Vector Architecture (SEVA) format^36^. Plasmids were constructed either using In-Fusion Snap Assembly (Takara Bio), unless otherwise noted, and verified by whole-plasmid Nanopore sequencing. A complete list of plasmids, assembly primers, and notes can be found in Supplementary Table 5. Base strains carrying poly-*attB* landing pads were generated for each PNSB host to enable site-specific integration. The *R. sphaeroides* landing pad and restriction endonuclease gene deletion strains were generated by two-step allelic exchange using pK18mobSacB-derived vectors. *R. palustris* P4, *R. rubrum*, and *R. gelatinosus* landing pads were delivered by a Tn7 transposon-based system we adapted for this study, where a Tn7 delivery vector carrying the poly-*attB* cassette and flanking FRT sites allows for subsequent excision of the antibiotic resistance marker. The Tn7 expression plasmid was co-transformed as either a non-replicating (pJMP1039) or replicating (pJMP8045) helper plasmid. pJMP1039 was a gift from Carol Gross & Jason Peters & Oren Rosenberg (Addgene plasmid #119239; http://n2t.net/addgene:119239; RRID:Addgene_119239). pJMP8045 was a gift from Jason Peters (Addgene plasmid #220900; http://n2t.net/addgene:220900; RRID:Addgene_220900). We used PCR to verify correct cassette insertion into each organism. Reporter and pathway integration strains were generated by tri-parental conjugation using *E. coli* WM6026^42^ donors carrying the integrase expression plasmid and the cognate *attP* payload plasmid. Single colonies were screened for correct integration by colony PCR and reporter fluorescence, and confirmed integrants were archived for subsequent engineering steps. For sequential integration experiments, orthogonal serine integrases (Bxb1, TG1, and R4) were paired with distinct antibiotic resistance markers and reporters (sfGFP, mRFP, and mTagBFP2) to enable iterative selection without marker curing, as described in the section “Sequential integration of multiple payloads”. Marker recycling was performed using ϕC31 integrase-mediated excision and *sacB* counterselection, as described in the section “Marker recycling via ϕC31 integrase and *sacB* counterselection”. To facilitate community adoption of cSAGE, we have deposited a curated set of plasmids with Addgene, summarized in Table 2.

**Table 2.**
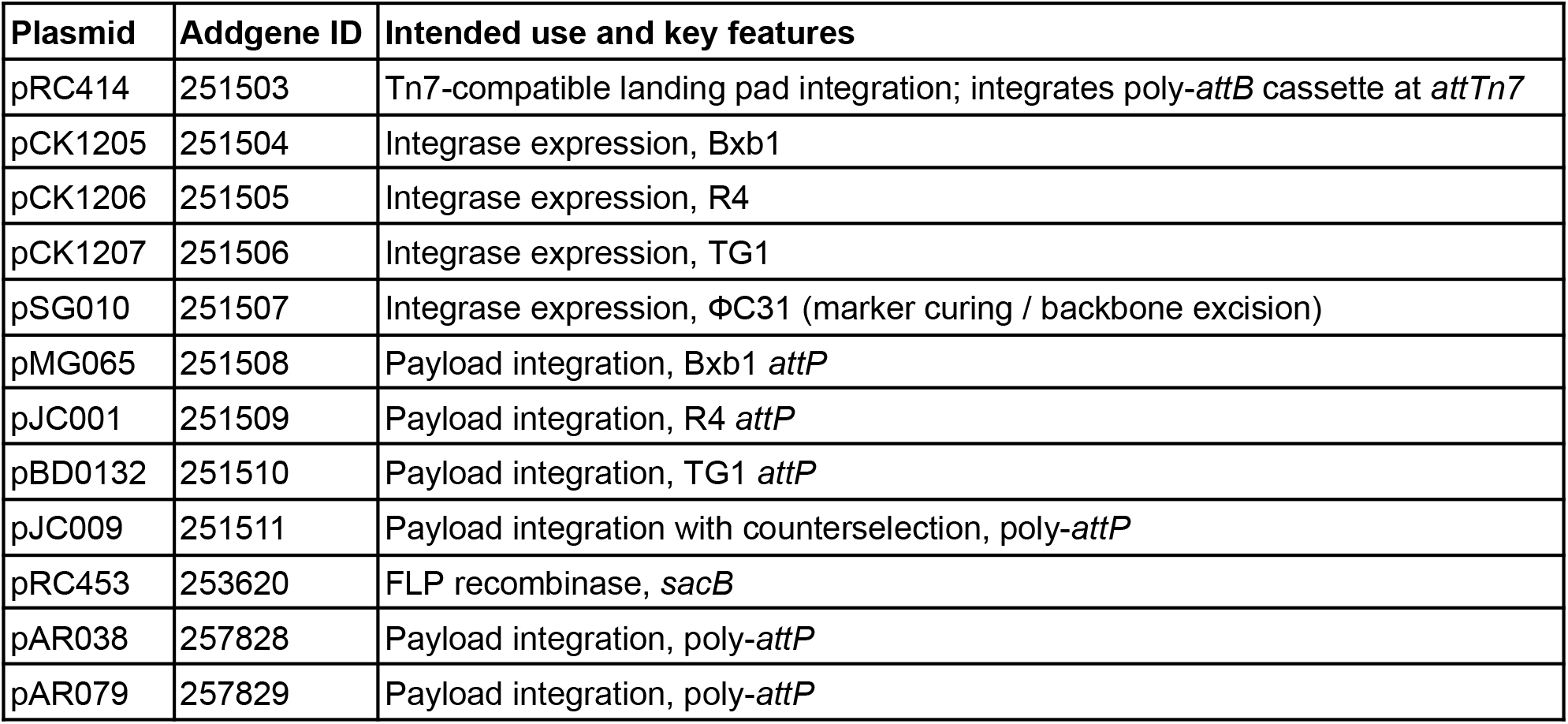
cSAGE plasmid toolkit.

| Plasmid | Addgene ID | Intended use and key features |
| --- | --- | --- |
| pRC414 | 251503 | Tn7-compatible landing pad integration; integrates poly-*attB* cassette at *attTn7* |
| pCK1205 | 251504 | Integrase expression, Bxb1 |
| pCK1206 | 251505 | Integrase expression, R4 |
| pCK1207 | 251506 | Integrase expression, TG1 |
| pSG010 | 251507 | Integrase expression, ΦC31 (marker curing / backbone excision) |
| pMG065 | 251508 | Payload integration, Bxb1 *attP* |
| pJC001 | 251509 | Payload integration, R4 *attP* |
| pBD0132 | 251510 | Payload integration, TG1 *attP* |
| pJC009 | 251511 | Payload integration with counterselection, poly-*attP* |
| pRC453 | 253620 | FLP recombinase, *sacB* |
| pAR038 | 257828 | Payload integration, poly-*attP* |
| pAR079 | 257829 | Payload integration, poly-*attP* |

**Table 3.** Comparison of cSAGE to alternative genome engineering approaches.

| **Feature** | **Tn7-based systems** | **CRAGE** | **CRISPR/HDR** | **SAGE** | **cSAGE** |
| --- | --- | --- | --- | --- | --- |
| *Conjugative delivery* | ✓<sup>19,23</sup> | ✓<sup>25</sup> | ✓<sup>20</sup> | ✗<sup>26</sup> | ✓ (Fig. 2, 4) |
| *Iterative integrations* | ✗ (single site)<sup>19,23</sup> | ✗ capped at 2 sites (CRAGE-DUET)<sup>24</sup> | ✓ unlimited (new construct required per edit)<sup>20</sup> | ✓ (poly-*attB* landing pad)<sup>26</sup> | ✓ (poly-*attB* landing pad, Fig. 4) |
| *Marker recycling* | ✓ Flp-FRT<sup>19,23</sup> | ✓ Cre-*lox* cassette exchange<sup>24,25</sup> | ✓<sup>20</sup> | ✓ (φC31 excision)<sup>26</sup> | ✓ (φC31 excision, conjugation-compatible, Fig. 5) |
| *Standardized toolkit across hosts* | ✗ (*attTn7* only conserved in Gram-negatives)<sup>19,23</sup> | ✓ (validated across multiple phyla)<sup>25</sup> | ✗ (strain-specific HDR construct required)<sup>20</sup> | ✓ (validated across multiple phyla)<sup>26</sup> | ✓ (identical plasmids across two phyla; Table 1, Figs. 2-3) |

### Electrocompetent cell preparation

Electrocompetent cells were prepared as previously described for *R. palustris*^26^, with the same protocol applied to *R. sphaeroides* which is comparable to optimized approaches developed for this species^39^. Briefly, cells were grown aerobically to mid-to-late log phase (OD_600_ 0.5-0.9), harvested, and washed three times in ice-cold 10% glycerol before resuspension at 1:400 the original culture volume. Electroporation was performed in 0.1-cm cuvettes with ∼100 ng plasmid DNA for both the *attP*/cargo plasmid and serine recombinase plasmid at 1.75 kV, 25 μF, and 200 Ω. Cells were recovered for 6 h at 30°C with shaking prior to plating on selective medium.

### Serine recombinase screening assays

To assess integrase performance across hosts, we used a standard tri-parental conjugation protocol to deliver both the non-replicating poly-*attP* target plasmid and the corresponding non-replicating integrase expression plasmid into recipient strains (Supplementary Table 2). All conjugations were carried out using the *E. coli* donor strain WM6026, which carries the *pir* genes necessary for R6K replication and is auxotrophic for diaminopimelic acid (DAP) to allow for counterselection post-mating. Donor WM6026 cells were grown overnight in LB with 0.1 mM DAP and appropriate antibiotics. Separately, PNSB recipients were cultured in rich medium (e.g., 3xYPS-MOPS for *R. palustris*, LB for *R. sphaeroides*) for 2-5 days until mid- to late-log phase. For tri-parental conjugations, WM6026 carrying the integrase plasmid and WM6026 carrying the poly-*attP* target plasmid were each washed and mixed with the recipient strain at equal optical densities (OD₆₀₀ = 1.0) in a 1:1:1 ratio. The cell mixtures were centrifuged, resuspended in ∼50 µL of fresh media, and spotted onto non-selective agar plates supplemented with 0.1 mM DAP and incubated overnight (∼16 h) to allow for conjugative DNA transfer and recombination. After overnight incubation at 30°C, conjugation mixtures were harvested, washed three times in fresh medium to remove residual DAP, and serially diluted for plating onto selective agar (lacking DAP) containing the appropriate antibiotic for selection of a single-*attP* or poly-*attP* target plasmid. Control recipient-only plates were used to quantify conjugation efficiency as transconjugants per viable recipient. Colonies were counted after 3–5 days of incubation at 30°C. Integration accuracy was assessed by colony PCR using a primer flanking the intended *attB* site and a single primer internal to the integrated target plasmid (Supplementary Table 5, Supplementary Fig. 10). In *R. palustris* CGA009 and P4, *R. rubrum*, and *R. gelatinosus*, kanamycin selection (poly-*attP* plasmid pSA29) was used to quantify conjugation efficiency following delivery of the payload and recombinase plasmids. However, in *R. sphaeroides*, kanamycin selection proved unreliable, likely due to antibiotic marker incompatibility, so we substituted spectinomycin (poly-*attP* plasmid pAR038) as the selectable marker unless otherwise indicated (Supplementary Fig. 4). Conjugation efficiencies were determined as the number of colonies on selective media (transconjugants) per number of viable recipient cells.

### Tn7 benchmarking study

To benchmark cSAGE against a standard Tn7-based delivery approach under matched conditions, we performed tri-parental conjugation using the mini-Tn7 helper plasmid pJMP1039 (R6K-based Tn7 transposase) and the *attTn7* cargo plasmid pCK383 (J23111::mRFP, spectinomycin resistance marker) (Supplementary Table 2), following the tri-parental mating protocol described above for recombinase screening, with *R. sphaeroides* as the recipient. Mating mixtures were plated on selective medium (spectinomycin) across a dilution series (10^−1^ to 10^−7^), and viable recipient cell density was determined in parallel by plating the same OD-normalized recipient culture on non-selective medium. No transconjugant colonies were recovered at any dilution across two independent conjugations, yielding a calculated detection limit of ≤10-10 transconjugants per recipient CFU (Supplementary Fig. 2).

### Reporter fluorescence quantification

Single colonies were inoculated in a 2 ml 96-DWP containing rich broth media with antibiotic selection (500 µL media). Plates were incubated at 30°C on a plate shaker for 24-48 h to enable sufficient outgrowth and fluorescent reporter expression. Optical density (OD_660_) and fluorescence were measured on a BioTek Synergy H1 plate reader and compared to WT controls to ascertain RFU/OD_660_. The following plate reader settings were used, unless otherwise noted: sfGFP: 485nm/528nm, mTagBFP2: 400nm/455nm, and mRFP 540nm/600nm.

### Promoter activity assays

To benchmark promoter activity in *R. palustris* CGA009 and *R. sphaeroides*, Anderson series (https://parts.igem.org/) promoters were cloned upstream of an mRFP reporter and integrated into the genome at the SAGE landing pad (Supplementary Tables 2 and 3). Strains were cultivated under routine aerobic cultivation conditions at 30°C, and fluorescence was measured (Ex/Em: 540nm/600nm) and calculated as described above.

### Marker recycling via ϕC31 integrase and sacB counterselection

To enable antibiotic marker recycling following integration, we adapted a marker curing strategy using ϕC31 integrase-mediated excision and *sacB*-based counterselection (Elmore et al. 2023). PNSB strains containing a genomically integrated poly-*attP* target plasmid (pMG97 or pJC009) were used as recipients for the ϕC31 plasmid (pSG010). WM6026 donor strains harboring the non-replicating ϕC31 integrase expression plasmid were prepared as described above and conjugated into these strains using di-parental mating. After mating, cells were plated onto agar containing sucrose (concentrations used for each strain are noted in Supplementary Table 4), excluding DAP, to select for marker loss and counterselect *E. coli*. Purple colonies appearing after 3–5 days of incubation at 30°C were screened by colony PCR using primers flanking the Bxb1 *attB* site (Supplementary Table 5). Successful marker curing was indicated by the loss of the antibiotic marker and plasmid backbone, while retention of the payload was verified by *mRFP* fluorescence.

### Sequential integration of multiple payloads

To demonstrate multi-payload genome engineering using cSAGE, we constructed a three-plasmid toolkit in which each plasmid encodes a different fluorescent reporter (sfGFP, mTagBFP2, or mRFP), a unique integrase attachment site (Bxb1, R4, or TG1), and an orthogonal antibiotic resistance marker (Km^R^, Sm^R^, or Gm^R^). Integration was performed sequentially in *R. palustris* CGA009 and *R. sphaeroides* using tri-parental conjugation as described above, starting first with mTagBFP2 (pJC001), then mRFP (pMG083 for CGA009 or pJC008 for *R. sphaeroides*), and lastly, sfGFP (pBD0132). After each integration round, transconjugants were screened for reporter expression and validated by colony PCR to confirm integration at each intended *attB* site (Supplementary Table 5). Confirmed single-copy integrants were subcultured and used as recipients for the subsequent integration. Each round of integration was performed using only the antibiotic marker relevant to the newly introduced plasmid, with no selection for previously integrated markers. Fluorescence was quantified for each strain to evaluate reporter expression and confirm stable maintenance of all three integrated payloads.

### Genomic integration of BaPAD

*p*-VP producing strains were generated by tri-parental conjugation of pRC442 (J23110::BaPAD) and pCK1205 (Bxb1 helper plasmid) with either *R. palustris* CGA009 (JE4632), *R. palustris* P4 (sMG127) or *R. sphaeroides* 2.4.1 (AVS4) (Supplementary Table 1). pRC442 contains a codon-optimized phenolic acid decarboxylase gene from *Bacillus atrophaeus* (BaPAD), the Bxb1 *attP* site, and a kanamycin selection marker. Successful integrants were screened as described above.

### Bioproduction of p-vinylphenol

Metabolites were quantified by preparing a standard curve in Sistrom’s media and fitting it to a linear equation. Media containing *p*-coumarate (*p*-CA) was prepared as previously described^80^. As a result of *p*-CA insolubility, sonication followed by filtration was used to make 10 mM *p*-CA in Sistrom’s minimal media. *R. palustris* strains were fed exclusively *p*-CA. Subsequent bioproduction experiments were inoculated from a starting anaerobic culture. Cultures were incubated at 30°C with infrared light until reaching stationary phase (7 days), at which point they were sampled periodically. For quantitative analysis, 1 mL of cell cultures were extracted using a 1 mL syringe and centrifuged for 5 min at 20,000×g. Supernatants were loaded into 0.2 μm, 10 kDa PES polypropylene filter 96-well plates (Analytical, cat. no. 964014) and centrifuged for 20 min at 1,500×g. *p*-vinylphenol (*p*-VP) and *p*-CA were quantified using high-performance liquid chromatography (HPLC). Filtered supernatants were collected in HPLC vials with piercable caps. Samples were analyzed by injecting 50 μL into an Agilent HPLC system equipped with a diode array detector set at 254 nm and 310 nm and separated using a ZORBAX RX-C18 reverse-phase column (4.6 nm × 150 nm, 5 μm particle size; Agilent Technologies). The mobile phases were (A) acetonitrile plus 0.1% trifluoroacetic acid (TFA), and (B) water plus 0.1% TFA. The flow was held at 1.000 mL/min. The mobile phase gradient follows: hold at 10% solvent A from 0.0 to 2.0 min, 10% to 50% solvent A from 2.0 to 12.0 min, hold at 50% solvent A from 12.0 to 14.0 min, 50% to 10% solvent A from 14.0 to 14.5 min, hold at 10% solvent A from 14.5 to 20.0 min. The retention times for the detected compounds were: 11.5 min (*p*-VP) and 7.1 min (*p*-CA). *p*-CA and *p*-VP concentrations were determined according to a standard curve of known concentrations (Supplementary Fig. 9).

### Optical density measurements

Optical density (OD_660_) of bacterial cultures in Hungate tubes was measured using a custom-built, non-invasive photometer adapted from the TubeOD design^81^. The device consists of a 3D-printed tube holder with an LED (596 nm) and a photosensor positioned on opposite sides of the culture tube to measure transmitted light (Supplementary Fig. 11). Light intensity was recorded as an analog voltage inversely proportional to cell density and logged using a LabJack U3-HV. Species-specific calibration curves were generated relative to a benchtop spectrophotometer and used to convert voltage to OD_660_ values. Python scripts are located on the GitHub repository https://github.com/carothersresearch/K-BOT.

### Genome-scale metabolic modeling

Flux values for the metabolic map of *R. palustris* CGA009 were calculated using COBRApy with a genome-scale model based on iAN1128, with additional reactions to represent the heterologous PAD enzyme and transport reactions for the product of the engineered pathway, *p*-vinylphenol. Simulations were performed under photoheterotrophic conditions with *p*-CA as the sole carbon source. The code used to generate these flux values is available at https://github.com/carothersresearch/cSAGE.

### Comparative genomic analysis

Alignments of the coumarate catabolic pathway genes between *R. palustris* CGA009 and P4 were performed using Mauve plugin in Geneious Prime v2026.0. We sequenced and assembled the P4 genome using Oxford Nanopore Sequencing serviced by Plasmidsaurus Inc. (with 100x coverage depth) and deposited it to GenBank under accession number CM151374.1. KEGG orthology (KO) analysis was performed using BLASTKoala for KO assignment. Restriction endonucleases were identified in the *R. sphaeroides* 2.4.1 genome assembly (GenBank accession number GCF_000012905) by scanning for the Gene Ontology (GO) term for endonuclease activity (GO:0004519). *RshI* was previously characterized^38^.

### Statistics and Reproducibility

Replicate numbers and experimental units are stated in each figure legend. Experiments used independent biological cultures or independent conjugation reactions as specified. For fluorescence and growth assays, values represent the mean of independent cultures with error bars indicating one standard deviation. For HPLC-based metabolite quantification, standard curves were prepared fresh for each experiment, and quantification was performed using at least three biological replicates. Integration efficiencies are reported as the mean proportion of colonies with correct integration, as determined by colony PCR.

## Data availability

All data in this study are available from the corresponding authors upon request. The source data underlying Figs. 1c-d, 2c, 3, 4b-c, 6b-c and Supplementary Figs. 3-5, 7, 9, and 11 are provided as a Source Data file. Key plasmids from the cSAGE toolkit have been deposited to Addgene (accession numbers listed in Table 2). Additional plasmids generated in this study are available from the corresponding authors upon request.

## Author contributions

†M.S.G. and C.K. contributed equally to this work. ^+^J.C. and A.R. contributed equally to this work. M.S.G., C.K., R.L.C., D.A.B. and J.E. conceived and designed the study. M.S.G., C.K., J.C., A.R., A.S., R.L.C., D.A.B., M.C., S.G., S.A., B.D., G.N.D, K.H. and D.H. performed experiments and analyzed the data. M.S.G., J.M.C., J.Z., J.E., R.G.E., A.M.G., and A.B. supervised the project and guided data interpretation. A.B. and J.M.C. acquired funding. M.S.G., C.K., A.R., J.C., R.L.C., M.C., J.E., and J.M.C. wrote the manuscript with input from all authors.

## Acknowledgements

We thank: Carrie Harwood and Yasuhiro Oda at the University of Washington for providing essential strains, protocols, and reagents that enabled this work; Huimin Zhao at the University of Illinois Urbana-Champaign for sharing *E. coli* WM6026; Eric Schaedig at The National Renewable Energy Laboratory and Carrie Eckhert at Oak Ridge National Laboratory who provided *Rubrivivax gelatinosus* CBS, as well as guidance and protocols for working with this strain. This research was supported by the DOE Office of Science, Office of Biological and Environmental Research (BER), grant no. DE-SC0023091. This report was prepared as an account of work sponsored by an agency of the United States Government. Neither the United States Government nor any agency thereof, nor any of their employees, makes any warranty, express or implied, or assumes any legal liability or responsibility for the accuracy, completeness, or usefulness of any information, apparatus, product, or process disclosed, or represents that its use would not infringe privately owned rights. Reference herein to any specific commercial product, process, or service by trade name, trademark, manufacturer, or otherwise does not necessarily constitute or imply its endorsement, recommendation, or favoring by the United States Government or any agency thereof. The views and opinions of the authors expressed herein do not necessarily state or reflect those of the United States Government or any agency thereof.

