## Supplementary information for "Conjugative serine recombinase engineering enables genome integration in transformation-resistant bacteria and bioproduction pathway prototyping"

**Supplemental Information**

Conjugative serine recombinase engineering extends high-efficiency genome integration and bioproduction to transformation-resistant bacteria

Michael S. Guzman[1,2*†], Cholpisit Kiattisewee[1,2,†], Amanda M. Robert[1,2+], Jackson Comes[1,2+], Ryan A.L. Cardiff[1,2], Allan Scott[3], Margaret Cook[1,2], Diego Alba Burbano[1,2], Stella Anastasakis[1,2], Sarah Grube[1,2], Kieran Heiberg[1], Brian Darst[1,2], Daniel Howell[1,2], Gara N. Dexter[4], Robert G. Egbert[3], Jesse G. Zalatan[1,2,5], Adam M. Guss[4], Joshua R. Elmore[3,*], Alexander S. Beliaev[6,*], James M. Carothers[1,2*]

1: Department of Chemical Engineering
University of Washington
Seattle, WA 98195
United States

2: UW Center for Synthetic Biology and Molecular Engineering & Sciences Institute
University of Washington
Seattle, WA 98195
United States

3: Biological Sciences Division, Pacific Northwest National Laboratory
Richland, WA 99354
United States

4: Biosciences Division, Oak Ridge National Laboratory
Oak Ridge, TN 37831
United States

5: Department of Chemistry
University of Washington
Seattle, WA 98195
United States

6: Environmental Molecular Sciences Division,
Pacific Northwest National Laboratory
Richland, WA 99354
United States

* Correspondence to:
Michael S. Guzman:
Joshua Elmore:
Alex Beliaev:
James M. Carothers:

**Supplementary Table 1.** Bacterial strains used in this study.

| Strain | Genotype and use | Source |
|---|---|---|
| *Cloning* | | |
| WM6026 | [lacIq rrnB3 ΔlacZ4787 hsdR514ΔaraBAD567 ΔrhaBA D568 rph-1 attl::pAE12(ΔoriR6K-cat::Frt5) ΔendA::Frt uidA(ΔMluI)::pir attHK::pJK1006Δ(oriR6K-cat::Frt5;trfA::Frt)]. Donor strain for conjugation. | Huimin Zhao, UIUC |
| EC100D *pir+* | *F - mcrA Δ(mrr-hsdRMS-mcrBC) φ80dlacZΔM15 ΔlacX74 recA1 endA1 araD139 Δ(ara, leu)7697 galU galK λ- rpsL (Str R) nupG pir +(DHFR)* | Fisher Scientific |
| NEB® Turbo | F' *proA+B+ lacIq ∆lacZM15 / fhuA2 ∆(lac-proAB) glnV galK16 galE15 R(zgb-210::Tn10)*TetS *endA1 thi-1 ∆(hsdS-mcrB)5* | NEB |
| *Wild-type* | | |
| CGA009 | *Rhodopseudomonas palustris* CGA009 | Carrie Harwood, UW |
| 2.4.1 | *Rhodobacter sphaeroides* 2.4.1 | Joshua Elmore, PNNL |
| CBS | *Rubrivivax gelatinosus* CBS | NREL |
| 11170 | *Rhodospirillum rubrum* ATCC 11170 | ATCC |
| P4 | *Rhodopseudomonas palustris* P4 | Joshua Elmore, PNNL |
| KT2440 | *Pseudomonas putida* KT2440 | Joshua Elmore, PNNL |
| RHA1 | *Rhodococcus jostii* RHA1 | Joshua Elmore, PNNL |
| SO81 | *Pseudomonas sorgoleonovorans* SO81 | Joshua Elmore, PNNL |
| *SAGE landing pad strains* | | |
| AVS2 | *R. sphaeroides* 2.4.1 Δ*RshI::[*10xpoly-*attB* ] | This study |
| AVS3 | *R. sphaeroides* 2.4.1 Δ*RshI::[*10xpoly-*attB* ]; Δ*RSP_RS17455* | This study |
| AVS4 | *R. sphaeroides* 2.4.1 Δ*RshI::[*10xpoly-*attB* ]; Δ*RSP_RS17455*; Δ*RSP_RS21745* | This study |
| JE4632 | *R. palustris* CGA009 Δ*RPA1300::[10xpoly-attB]* | Elmore *et al.* 2023 |
| sMG127 | *R. palustris* P4 *attTn7::*[10×poly-*attB*] | This study |
| sMG223 | *R. rubrum* ATCC 11170 *attTn7::*[10×poly-*attB*] | This study |
| sMG229 | *R. gelatinosus* CBS *attTn7::*[10×poly-attB]*::GmR* | This study |
| KT2440 | *Pseudomonas putida* KT2440 [10×poly-*attB*] | This study |
| RHA1 | *Rhodococcus jostii* RHA1 [10×poly-*attB*] | This study |
| SO81 | *Pseudomonas sorgoleonovorans* SO81 [10×poly-*attB*] | This study |
| *p-VP production* | | |
| sRC050 | sMG127 *attB_Bxb1*::pRC442 | This study |
| sSA015 | JE4632 *attB_Bxb1*::pRC442 | This study |
| sAR167 | AVS4 *attB_Bxb1*::pRC442 | This study |

**Supplementary Table 2.** Plasmids used in this study.

| Plasmid | Genotype | Intended Use | Source |
|---|---|---|---|
| *Integrases* | | | |
| pCK1205 | $P_{tac}$::*Bxb1*; $Amp^R$; $ori_{R6K}$ | Constitutive integrase | This study |
| pCK1206 | $P_{tac}$::*R4*; $Amp^R$; $ori_{R6K}$ | Constitutive integrase | This study |
| pCK1207 | $P_{tac}$::*TG1*; $Amp^R$; $ori_{R6K}$ | Constitutive integrase | This study |
| pDH001 | $P_{tac}$::*ΦBT1*; $Amp^R$; $ori_{R6K}$ | Constitutive integrase | This study |
| pDH002 | $P_{tac}$::*RV*; $Amp^R$; $ori_{R6K}$ | Constitutive integrase | This study |
| pSG007 | $P_{tac}$::*MR11*; $Amp^R$; $ori_{R6K}$ | Constitutive integrase | This study |
| pSG009 | $P_{tac}$::*ΦC1*; $Amp^R$; $ori_{R6K}$ | Constitutive integrase | This study |
| pSG010 | $P_{tac}$::*ΦC31*; $Amp^R$; $ori_{R6K}$ | Marker curing | This study |
| pSG011 | $P_{tac}$::*BL3*; $Amp^R$; $ori_{R6K}$ | Constitutive integrase | This study |
| *Reporters* | | | |
| pCK1137 | J23119::*sfGFP*; $Km^R$; $ori_{R6K}$; $attP_{Bxb1}$ | Sequential integrations | This study |
| pBD0131 | J23119::*sfGFP*; $Sm^R$; $ori_{R6K}$; $attP_{R4}$ | Sequential integrations | This study |
| pBD0132 | J23119::*sfGFP*; $Gm^R$; $ori_{R6K}$; $attP_{TG1}$ | Sequential integrations | This study |
| pSA029 | J23108::*mRFP*; $Km^R$; $ori_{R6K}$; $attP_{poly}$ | Integrase screening | This study |
| pMG083 | J23119::*sfGFP*; *sacB*; $Km^R$; $ori_{R6K}$; $attP_{Bxb1}$ | Marker curing | This study |
| pJC001 | J23119::*mTagBFP2*; $Sm^R$; $ori_{R6K}$; $attP_{R4}$ | Sequential integrations | This study |
| pJC008 | J23119::*mRFP*; $Km^R$; $ori_{R6K}$; $attP_{Bxb1}$ | Sequential integrations | This study |
| pJC009 | J23108::*mRFP*; *sacB*; $Km^R$-pBBR1MCS-2; $ori_{R6K}$; $attP_{poly}$ | Marker curing | This study |
| pMG097 | J23108::*mRFP*; *sacB*; $Km^R$; $ori_{R6K}$; $attP_{poly}$ | Marker curing | This study |
| pMG053 | J23109::*mRFP*; $Km^R$; $ori_{R6K}$; $attP_{Bxb1}$ | Promoter screening | This study |
| pMG055 | J23113::*mRFP*; $Km^R$; $ori_{R6K}$; $attP_{Bxb1}$ | Promoter screening | This study |
| pMG056 | J23117::*mRFP*; $Km^R$; $ori_{R6K}$; $attP_{Bxb1}$ | Promoter screening | This study |
| pMG057 | J23114::*mRFP*; $Km^R$; $ori_{R6K}$; $attP_{Bxb1}$ | Promoter screening | This study |
| pMG058 | J23115::*mRFP*; $Km^R$; $ori_{R6K}$; $attP_{Bxb1}$ | Promoter screening | This study |
| pMG059 | J23107::*mRFP*; $Km^R$; $ori_{R6K}$; $attP_{Bxb1}$ | Promoter screening | This study |
| pMG060 | J23105::*mRFP*; $Km^R$; $ori_{R6K}$; $attP_{Bxb1}$ | Promoter screening | This study |
| pMG061 | J23106::*mRFP*; $Km^R$; $ori_{R6K}$; $attP_{Bxb1}$ | Promoter screening | This study |
| pMG062 | J23108::*mRFP*; $Km^R$; $ori_{R6K}$; $attP_{Bxb1}$ | Promoter screening | This study |
| pMG063 | J23110::*mRFP*; $Km^R$; $ori_{R6K}$; $attP_{Bxb1}$ | Promoter screening | This study |
| pMG064 | J23111::*mRFP*; $Km^R$; $ori_{R6K}$; $attP_{Bxb1}$ | Promoter screening | This study |
| pMG065 | J23119::*mRFP*; $Km^R$; $ori_{R6K}$; $attP_{Bxb1}$ | Promoter screening | This study |
| pAR038 | J23108::*mRFP*; $Sm^R$; $ori_{R6K}$; $attP_{poly}$ | Integrase screening | This study |
| pAR039 | J23108::*mRFP*; $Sm^R$; $ori_{R6K}$; $attP_{Bxb1}$ | Promoter screening | This study |
| pAR050 | J23105::*mRFP*; $Sm^R$; $ori_{R6K}$; $attP_{Bxb1}$ | Promoter screening | This study |
| pAR051 | J23106::*mRFP*; $Sm^R$; $ori_{R6K}$; $attP_{poly}$ | Integrase screening | This study |
| pAR052 | J23111::*mRFP*; $Sm^R$; $ori_{R6K}$; $attP_{Bxb1}$ | Promoter screening | This study |
| pAR053 | J23119::*mRFP*; $Sm^R$; $ori_{R6K}$; $attP_{Bxb1}$ | Promoter screening | This study |
| pAR054 | J23108::*mRFP*; $Km^R$-pBBR1MCS-2; $ori_{R6K}$; $attP_{poly}$ | Integrase screening | This study |
| pCK421 | J23106::*sfGFP,* $Km^R$; $ori_{RK2}$ | Marker screening | This study |
| pCK524 | J23106::*sfGFP,* $Km^R$; $ori_{BBR1}$ | Marker screening | This study |
| pCK564 | J23106::*sfGFP,* $Km^R$; $ori_{RSF1010}$ | Marker screening | This study |

| | | | |
|---|---|---|---|
| *Knockout/in* | | | |
| pJQ200SK | SacB; $Gm^R$; $ori_{p15A}$; $oriT_{RP4}$ | Base allelic exchange vector | Harwood Lab, UW |
| pAVS2 | pJQ200SK Δ*RshI*::[10xpoly-*attB*] | Allelic exchange vector | This study |
| pAVS3 | pJQ200SK Δ*RSP_RS17455* | Allelic exchange vector | This study |
| pAVS4 | pJQ200SK Δ*RSP_RS21745* | Allelic exchange vector | This study |
| *Tn7* | | | |
| pJMP1039 | Tn7 transposase (pR6K); $Amp^R$ | Tn7 integration | Addgene |
| pJMP8045 | Tn7 transposase (pBBR1); $Sm^R$ | Tn7 integration | Addgene |
| pRC453 | Flp recombinase; SacB; $Sm^R$; $ori_{BBR1}$ | Tn7 integration | This study |
| pRC414.v2 | attTn7::[10xpoly-*attB*]; $Amp^R$; $Gm^R$; $ori_{R6K}$ | Tn7 integration | This study |
| pCK383 | attTn7::[J23111::*mRFP*]; $Amp^R$; $Sm^R$; $ori_{R6K}$ | Tn7 integration | This study |
| *Replicating vectors* | | | |
| pCK205 | J23109::*mRFP,* $Gm^R$; $ori_{pBBR1}$ | Promoter screening | This study |
| pCK184 | J23113::*mRFP,* $Gm^R$; $ori_{pBBR1}$ | Promoter screening | This study |
| pCK120 | J23117::*mRFP,* $Gm^R$; $ori_{pBBR1}$ | Promoter screening | This study |
| pCK206 | J23114::*mRFP,* $Gm^R$; $ori_{pBBR1}$ | Promoter screening | This study |
| pCK185 | J23115::*mRFP,* $Gm^R$; $ori_{pBBR1}$ | Promoter screening | This study |
| pCK186 | J23107::*mRFP,* $Gm^R$; $ori_{pBBR1}$ | Promoter screening | This study |
| pCK286 | J23105::*mRFP,* $Gm^R$; $ori_{pBBR1}$ | Promoter screening | This study |
| pCK287 | J23106::*mRFP,* $Gm^R$; $ori_{pBBR1}$ | Promoter screening | This study |
| pCK285 | J23108::*mRFP,* $Gm^R$; $ori_{pBBR1}$ | Promoter screening | This study |
| pCK207 | J23110::*mRFP,* $Gm^R$; $ori_{pBBR1}$ | Promoter screening | This study |
| pCK208 | J23111::*mRFP,* $Gm^R$; $ori_{pBBR1}$ | Promoter screening | This study |
| pCK391 | J23119::*mRFP,* $Gm^R$; $ori_{pBBR1}$ | Promoter screening | This study |
| pCK421 | J23106::*sfGFP,* $Km^R$; $ori_{pRK2}$ | Origin screening | This study |
| pCK564 | J23106::*sfGFP,* $Km^R$; $ori_{pRSF1010}$ | Origin screening | This study |
| pCK524 | J23106::*sfGFP,* $Km^R$; $ori_{pBBR1}$ | Origin screening | This study |
| *p-VP production* | | | |
| pRC442 | J23110::BaPAD, $Km^R$, $ori^{R6K}$, $attP_{Bxb1}$ | *p*-VP Production | This study |

**Supplementary Table 3.** Anderson promoter sequences used in this study (https://parts.igem.org/Promoters/Catalog/Anderson).

| Promoter | Sequence |
|---|---|
| BBa_J23109 | tttacagctagctcagtcctagggactgtgctagc |
| BBa_J23113 | ctgatggctagctcagtcctagggattatgctagc |
| BBa_J23117 | ttgacagctagctcagtcctagggattgtgctagc |
| BBa_J23114 | tttatggctagctcagtcctaggtacaatgctagc |
| BBa_J23115 | tttatagctagctcagcccttggtacaatgctagc |
| BBa_J23107 | tttacggctagctcagccctaggtattatgctagc |
| BBa_J23105 | tttacggctagctcagtcctaggtactatgctagc |
| BBa_J23106 | tttacggctagctcagtcctaggtatagtgctagc |
| BBa_J23108 | ctgacagctagctcagtcctaggtataatgctagc |
| BBa_J23110 | tttacggctagctcagtcctaggtacaatgctagc |
| BBa_J23111 | ttgacggctagctcagtcctaggtatagtgctagc |
| BBa_J23119 | ttgacagctagctcagtcctaggtataatactagt |

**Supplementary Table 4.** Antibiotics and counterselection conditions used for PNSB in this study.

| Species | Media | Kanamycin | Gentamicin | Spectinomycin | Sucrose |
|---|---|---|---|---|---|
| *R. sphaeroides* 2.4.1 | LB | 50 μg/ml | 60 μg/ml | 50 μg/ml | 10% |
| *R. palustris* CGA009 | 3xYPSMOPS | 200 μg/ml | 200 μg/ml | 200 μg/ml | 10% |
| *R. rubrum* ATCC 11170 | 3xYPSMOPS | 100 μg/ml | 30 μg/ml | 100 μg/ml | 10% |
| *R. gelatinosus* CBS | 3xYPSMOPS | 50 μg/ml | 30 μg/ml | 100 μg/ml | 1-10% |
| *R. palustris* P4 | 3xYPSMOPS | 50 μg/ml | 200 μg/ml | 100 μg/ml | 15% |

**Supplementary Table 5.** Primers used in this study for plasmid assembly and strain screening.

| Intended use | Primer | Sequence | Notes |
|---|---|---|---|
| *Plasmid assembly* | | | |
| pCK1205-1207 | oCK1843 | ctttttggttaactgcggccgcGTTTAAACcttggactcctgttg | AmpR-oriT-pR6K from pCK468 |
| | oCK1844 | atttgttttggtccactagtTTAATTAAaggcatcaaataaaacg | AmpR-oriT-pR6K from pCK468 |
| pDH001, pDH002, pSG007, pSG009-11 | oDH001 | gctagcatccaagtcttcaattg | pCK1205 backbone |
| | oDH002 | ctgcatcacctcctatgtgt | pCK1205 backbone |
| pDH001 | oDH003 | taggaggtgatgcagatgagcccgttcattgcg | ΦBT1 from pGW33 |
| | oDH004 | gacttggatgctagcttacagcgccgccagttc | ΦBT1 from pGW33 |
| pDH002 | oDH005 | taggaggtgatgcagatgcgttacaccacccc | RV from pGW32 |
| | oDH006 | gacttggatgctagcttaacgccagttcacttgaacac | RV from pGW32 |
| pSG007 | oSG024 | taggaggtgatgcagatgaaggttgcgatctacac | MR11 from pGW36 |
| | oSG025 | gacttggatgctagcttaatagaaatcgatgttcttgatcttg | MR11 from pGW36 |
| pSG009 | oSG034 | taggaggtgatgcagatgaaacgtgcggcgctgtac | ΦC1 from pGW34 |
| | oSG035 | gacttggatgctagcttaaaacttgtatttaatgataacctcatcg | ΦC1 from pGW34 |
| pSG010 | oSG038 | ataggaggtgatgcagatggacacctacgcgggc | ΦC31 from pJE1817 |
| | oSG039 | gacttggatgctagcttacgccgcaacatcctcgg | ΦC31 from pJE1817 |
| pSG011 | oSG040 | taggaggtgatgcagatgaaactgcgtgcggcg | BL3 from pGW40 |
| | oSG041 | gacttggatgctagcttagatgttccattcaatcttcatatcttc | BL3 from pGW40 |
| pCK1137 | oCK1809 | ttttcgttttggtccggatcTTGACAgctagctcagtcctaggTATAATgctagc | J23119-sfGFP-dblT from pCK464 |
| | oCK1810 | gccttattgttcgtcctcgaTATAAACGCAGAAAGGCCCACC | J23119-sfGFP-dblT from pCK464 |
| pSA029 | oSA077 | ggggctttctcatgcgttgctagc | poly-attP from pGW60 |
| | oSA080 | gacgaacaataaggcctccctaacggg | poly-attP from pGW60 |
| | oSA078 | cagttaaccaaaaagggggattttatctc | pMG061 backbone |
| | oSA079 | gcatgagaaagcccccgga | pMG061 backbone |
| pMG083 | oMG239 | gaatccaggggtcccctagaggatcgatccttttttaac | SacB from pAVS2 |
| | oMG240 | gacacaaatttaaatctcggtacccatcgg | SacB from pAVS2 |
| | oMG241 | ccgatgggtaccgagatttaaatttgtgtctcaaaatct | pCK1137 backbone |
| | oMG242 | ggatcgatcctctaggggacccctggattc | pCK1137 backbone |
| pJC001 | oCD242 | atgcgttatggtatcgatatgaacaccgactac | pCK1138 backbone |
| | oMG134 | tataaacgcagaaaggcccac | pCK1138 backbone |
| | oJC001 | gtgggcctttctgcgttta | mTagBFP2 from pMG010 |
| | oCDP004 | ggtacctttctcctctttaatgaattc | mTagBFP2 from pMG010 |
| pJC008 | oCK570 | gacttttgtccttttccgctgcataac | pMG065 backbone |
| | oAR106 | atttaaatcgtaattattggggacccctg | pMG065 backbone |
| | oAR108 | aattacgatttaaatgatgaatgtcagctactgggc | KmR (pBBR1) cassette from pCK565 |
| | oAR109 | gaaaaggacaaaagtcgggtgg | KmR (pBBR1) cassette from pCK565 |

| | | | |
|---|---|---|---|
| pJC009 | oCK570 | gacttttgtccttttccgctgcataac | pMG097 backbone |
| | oJC020 | atttaaatcgtaattattggggacccctgctcggtacccatcggcattttc | pMG097 backbone |
| | oAR108 | aattacgatttaaatgatgaatgtcagctactgggc | KmR (pBBR1) cassette from pCK565 |
| | oAR109 | gaaaaggacaaaagtcgggtgg | KmR (pBBR1) cassette from pCK565 |
| pMG097 | oMG245 | ctagaggatcgatcctttttaacccatcac | SacB from pAVS2 |
| | oMG247 | ctcggtacccatcggcattttcttttg | SacB from pAVS2 |
| | oMG241 | ccgatgggtaccgagatttaaatttgtgtctcaaaatct | pSA029 backbone |
| | oMG242 | ggatcgatcctctaggggacccctggattc | pSA029 backbone |
| pMG053-65 | oMG133 | AGCATTTGCGATCATTCACG | BBa_J23105-J23119::*mRFP* fragment from pCK205, pCK184, pCK120, pCK206, pCK185, pCK186, pCK286, pCK287, pCK285, pCK207, pCK208, pCK391 |
| | oMG134 | TATAAACGCAGAAAGGCCCAC | J3-J23XXX::mRFP from pCK205 |
| | oMG132 | ATGATCGCAAATGCTcggattgcaattgaagacttg | pCK1137 backbone |
| | oMG131 | CTTTCTGCGTTTATAataattcaggtagaagacaactgg | pCK1137 backbone |
| pAR038, pAR039, pAR050-53 | oAR080 | ccaaggtagtcggcaaataagacttttgtccttttccgctgc | pSA029 backbone |
| | oAR081 | ctgcgttcggtcaaggttcatttaaatcgtaattattggggaccc | pSA029 backbone |
| | oAR082 | atgaaccttgaccgaacgcag | SmR cassette from pBD0131 |
| | oAR083 | gtcttatttgccgactaccttgg | SmR cassette from pBD0131 |
| pAR054 | oAR080 | ccaaggtagtcggcaaataagacttttgtccttttccgctgc | pSA029 backbone |
| | oAR081 | ctgcgttcggtcaaggttcatttaaatcgtaattattggggaccc | pSA029 backbone |
| | oAR104 | agctcagtcctaggtatagtgctagcGAATTCATTAAAGAGGAGAAAGG | pAR038 backbone |
| | oAR105 | acctaggactgagctagccgtaaaTCTATAATCGCAACTTCAAGAC | pAR038 backbone |
| | oCK570 | gacttttgtccttttccgctgcataac | pAR051 backbone |
| | oAR106 | atttaaatcgtaattattggggacccctg | pAR051 backbone |
| | oAR108 | aattacgatttaaatgatgaatgtcagctactgggc | KmR (pBBR1) cassette from pCK565 |
| | oAR109 | gaaaaggacaaaagtcgggtgg | KmR (pBBR1) cassette from pCK565 |
| pAR079 | oAR142 | AGTTGCGATTATAGATTGACAgctagctcagtcctag | J23119::*sfGFP* From pMG083 |
| | oCK219 | GCCTGGagatccttactcga | J23119::*sfGFP* From pMG083 |
| | oMG067 | TCTATAATCGCAACTTCAAGACGACG | pJC009 backbone |
| | oCK064 | tcgagtaaggatctccaggc | pJC009 backbone |
| *Integration screening primers* | | | |
| | Bxb1_F | TGATCGAATTCTTCATTTAAAGACCCT | Elmore et al. 2023 |
| | Bxb1_R | GGCAGAATTTTGGGAGTGGGCAT | Elmore et al. 2023 |
| | BT1_F | TCCCAAATTCTGCCTAGAAAGTC | Elmore et al. 2023 |
| | BT1_R | TAGTGGAGGAACTAAAACAGC | Elmore et al. 2023 |
| | phiC1_F | GACTACGAGGCCGGATCA | Elmore et al. 2023 |
| | phiC1_R | TCCTCTTGAGGTAGAAACGGGG | Elmore et al. 2023 |
| | RV_F | GAATTCCGACATGGCAATAACCC | Elmore et al. 2023 |
| | RV_R | GCCGACAGCAGTCAGTTTATAC | Elmore et al. 2023 |

| | | | |
|---|---|---|---|
| | TG1_F | AAGTATTAACATGATCTGTCGG | Elmore et al. 2023 |
| | TG1_R | GGAGGATCCGATTGCATCCG | Elmore et al. 2023 |
| | R4_F | CGTCCACGGCGATGCAATG | Elmore et al. 2023 |
| | R4_R | GAAGTGTTCAACGCGTACGC | Elmore et al. 2023 |
| | BL3_F | TGGCGTACGCGTTGAACAC | Elmore et al. 2023 |
| | BL3_R | CACCCGCCTTTAGCTTCC | Elmore et al. 2023 |
| | A118_F | TAAGCGGGGTGAGAGGGTA | Elmore et al. 2023 |
| | A118_R | CCGTCAACATCAGTAGTCTCAC | Elmore et al. 2023 |
| | MR11_F | TGAGCATCTGATGTTGCACGG | Elmore et al. 2023 |
| | MR11_R | GAGTCGTGATCTGATAGTAGTGAG | Elmore et al. 2023 |
| | phi370_F | CGTTCGAGTATGAGTGCAGC | Elmore et al. 2023 |
| | phi370_R | AGCTGCAGAAGAACAATGCG | Elmore et al. 2023 |
| | oAR053 | CCGAGCGTTCTGAACAAATCC | Binds terminator of *attP* plasmids |

**Supplementary Table 6.** KEGG orthologies (KO) specific to *R. palustris* P4 based on BlastKOALA analysis.

| Functional Category | KO | Gene | Description |
|---|---|---|---|
| **Aromatic / Phenylpropanoid Metabolism** | K05710 | hcaC | Cinnamate / phenylpropionate dioxygenase component |
| **Transport / Efflux** | K01531 | mgtA/mgtB | $Mg^{2+}$ transporter |
| | K15551 | tauA | Taurine transporter (substrate-binding) |
| | K15552 | tauC | Taurine transporter (permease) |
| | K16092 | btuB | Vitamin B12 transporter |
| | K18902 | bpeF | Multidrug efflux pump |
| **Energy Metabolism / Electron Transport / Redox** | K00156 | poxB | Pyruvate dehydrogenase (quinone) |
| | K00198 | cooS/acsA | CO dehydrogenase |
| | K00244 | frdA | Succinate dehydrogenase |
| | K00428 | ccp | Cytochrome c peroxidase |
| | K02297 | cyoA | Cytochrome o oxidase subunit II |
| | K02298 | cyoB | Cytochrome o oxidase subunit I |
| | K02299 | cyoC | Cytochrome o oxidase subunit III |
| | K02300 | cyoD | Cytochrome o oxidase subunit IV |
| | K02569 | napC | Cytochrome c (Nap/Nir family) |
| | K07217 | ydbD | Manganese catalase |
| | K07321 | cooC | CO dehydrogenase maturation factor |
| | K11178 | yagS | Xanthine dehydrogenase (FAD subunit) |
| | K13483 | yagT | Xanthine dehydrogenase (Fe-S subunit) |
| | K13255 | fhuF | Ferric iron reductase |
| | K15408 | coxAC | Cytochrome c oxidase |
| | K17218 | sqr | Sulfide:quinone oxidoreductase |
| | K17230 | fccA | Sulfide dehydrogenase subunit |
| | K21636 | nrdD | Anaerobic ribonucleotide reductase |
| **Central Carbon Metabolism** | K01659 | prpC | 2-methylcitrate synthase |
| | K01720 | prpD | 2-methylcitrate dehydratase |
| | K04069 | pflA | Pyruvate formate lyase activating enzyme |
| **Regulation / Signaling** | K18144 | adeR | Two-component response regulator |
| | K21686 | prpR | Transcriptional regulator |
| **Sulfur Metabolism** | K01130 | atsA | Arylsulfatase |
| | K24118 | atsB | Sulfatase-maturating enzyme |
| **Amino Acid / Nitrogen Metabolism** | K01953 | asnB | Asparagine synthase |
| | K13049 | PM20D1 | Carboxypeptidase |
| **Cell Wall / Lysis** | K01185 | — | Lysozyme |
| **DNA Processing / Defense / Mobile Elements** | K01153 | hsdR | Type I restriction enzyme R |
| | K01154 | hsdS | Type I restriction enzyme S |
| | K03427 | hsdM | Type I restriction enzyme M |
| | K07481 | — | IS5 transposase |
| | K07491 | rayT | REP-associated transposase |
| | K06909 | xtmB | Phage terminase large subunit |

| | | | |
|---|---|---|---|
| | K07474 | xtmA | Phage terminase small subunit |
| | K07341 | doc | Toxin |
| | K18843 | hicB | Antitoxin |
| | K19092 | parE | Toxin |
| | K21495 | fitA | Antitoxin |
| **Stress / Redox / Cofactor** | K06048 | gshA | Glutathione biosynthesis |
| **Other / Unclassified** | K00526 | nrdB/nrdF | Ribonucleotide reductase beta |
| | K02351 | — | Putative membrane protein |
| | K06877 | — | DEAD/DEAH helicase |

**Supplementary Table 7.** KEGG orthologies (KO) specific to *R. palustris* CGA009 based on BlastKOALA analysis.

| Functional Category | KO | Gene | Description |
|---|---|---|---|
| **Aromatic / Phenylpropanoid Metabolism** | — | — | *(None identified)* |
| **Transport / Efflux** | K06188 | aqpZ/aqpM | Aquaporin |
| | K22736 | VIT | Vacuolar iron transporter |
| **Energy Metabolism / Electron Transport / Redox** | K00116 | mqo | Malate dehydrogenase (quinone) |
| | K02192 | bfd | Bacterioferritin-associated ferredoxin |
| | K07215 | pigA/hemO | Heme oxygenase |
| | K03741 | arsC | Arsenate reductase |
| | K07755 | AS3MT | Arsenite methyltransferase |
| | K11811 | arsH | Arsenical resistance protein |
| | K22896 | vnfD | Vanadium nitrogenase alpha chain |
| | K22897 | vnfK | Vanadium nitrogenase beta chain |
| | K22898 | vnfG | Vanadium nitrogenase delta subunit |
| | K22903 | vnfE | Nitrogenase cofactor synthesis protein |
| **Central Carbon / Lipid Metabolism** | K10254 | ohyA | Oleate hydratase |
| | K10255 | desA | Omega-6 fatty acid desaturase |
| | K14977 | ylbA | Ureidoglycine aminohydrolase |
| **Amino Acid / Nitrogen Metabolism** | K01487 | guaD | Guanine deaminase |
| | K01761 | — | Methionine gamma-lyase |
| **Regulation / Signaling** | K03719 | lrp | Leucine-responsive regulator |
| | K13652 | — | AraC family regulator |
| | K15735 | csiR | Carbon starvation regulator |
| | K13924 | cheBR | Chemotaxis regulator |
| | K18831 | higA | Transcriptional regulator / antitoxin |
| **Cell Surface / Extracellular Structures** | K16566 | exoY | Exopolysaccharide production protein |
| **DNA Processing / Repair / Epigenetics** | K03169 | topB | DNA topoisomerase III |
| | K07317 | — | DNA methyltransferase |
| | K07445 | — | DNA methylase |
| | K07458 | vsr | DNA mismatch repair endonuclease |
| **Secretion Systems / Conjugation** | K03205 | virD4 | Type IV secretion system protein |
| | K07344 | trbL | Type IV secretion system protein |
| | K20266 | trbJ | Type IV secretion system protein |
| | K20527 | trbB | Type IV secretion system protein |
| | K20528 | trbC | Type IV secretion system protein |
| | K20529 | trbD | Type IV secretion system protein |
| | K20530 | trbE | Type IV secretion system protein |
| | K20531 | trbF | Type IV secretion system protein |
| | K20532 | trbG | Type IV secretion system protein |
| | K20533 | trbI | Type IV secretion system protein |
| **Stress Response / Detoxification** | K03741 | arsC | Arsenate reductase |

| | | | |
|---|---|---|---|
| | K07755 | AS3MT | Arsenite methyltransferase |
| | K11811 | arsH | Arsenic resistance protein |
| **Mobile Elements / Phage / Toxin-Antitoxin** | K07486 | — | Transposase |
| | K07733 | alpA | Prophage regulator |
| | K19166 | higB | Toxin (mRNA interferase) |
| | K18831 | higA | Antitoxin |
| **Other / Unclassified** | K07089 | — | Uncharacterized protein |
| | K09138 | — | Uncharacterized protein |
| | K13525 | VCP/CDC48 | AAA ATPase |
| | K26273 | SBNO | Strawberry notch protein |

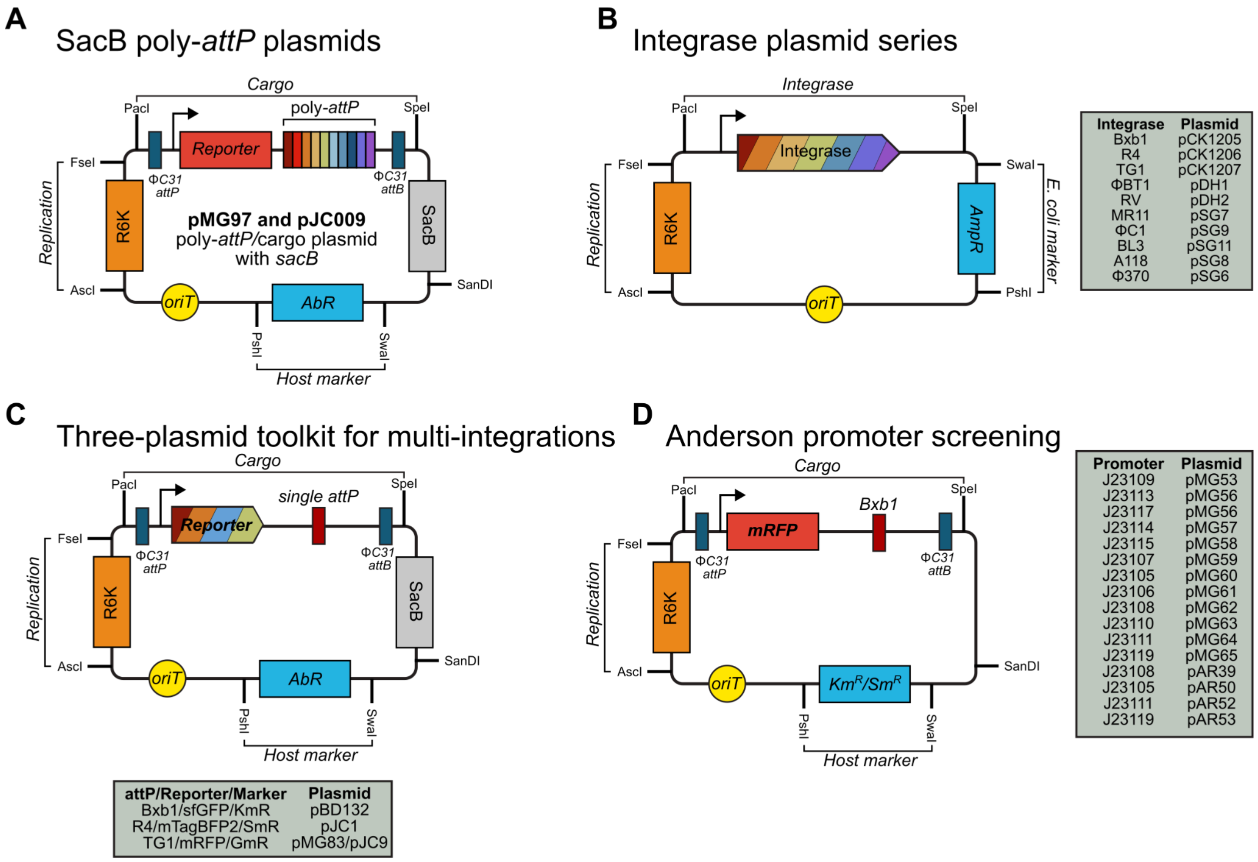


**Supplementary Figure 1.** The cSAGE modular plasmid toolkit in SEVA format. cSAGE plasmids were redesigned in the Standard European Vector Architecture (SEVA) format to enable modularity, portability, and compatibility with conjugation-based delivery. All vectors incorporate a *pir*-dependent R6K origin of replication to reduce metabolic burden in E. coli pir$^+$ donor strains and to prevent replication in recipient hosts, as well as an *oriT* for mobilization by conjugation. (A) Poly-*attP* plasmids for integration of payloads into SAGE landing pad. Each version carries a fluorescent reporter and a *sacB* cassette for sucrose-based counterselection to allow plasmid backbone curing after integration. Versions differ only in the antibiotic resistance marker for host selection. (B) Series of integrase helper plasmids, each encoding one of eight orthogonal serine recombinases. These plasmids provide transient integrase expression for site-specific recombination at cognate *attB*/*attP* sites. (C) Three-plasmid cSAGE toolkit enabling sequential multi-payload integration. Orthogonal *attP* cassettes (Bxb1, R4, TG1) were paired with distinct antibiotic resistance markers (Km$^R$, Sm$^R$, Gm$^R$) and fluorescent reporters (sfGFP, mTagBFP2, mRFP), supporting iterative rounds of integration without intermediate curing. (D) Library of Anderson promoters cloned upstream of mRFP with Bxb1 *attP* sites, used to benchmark promoter activity in *R. palustris* and *R. sphaeroides* at chromosomal landing pads. This plasmid series provides a resource for synthetic promoter characterization in PNSB (promoter sequences available at: https://parts.igem.org/Promoters/Catalog/Anderson).

**Recipient cells**
(*R. sphaeroides*)

**Tri-parental conjugation ($10^{-3}$)**
(pJMP1039 + pCK383 + *R. sphaeroides*)

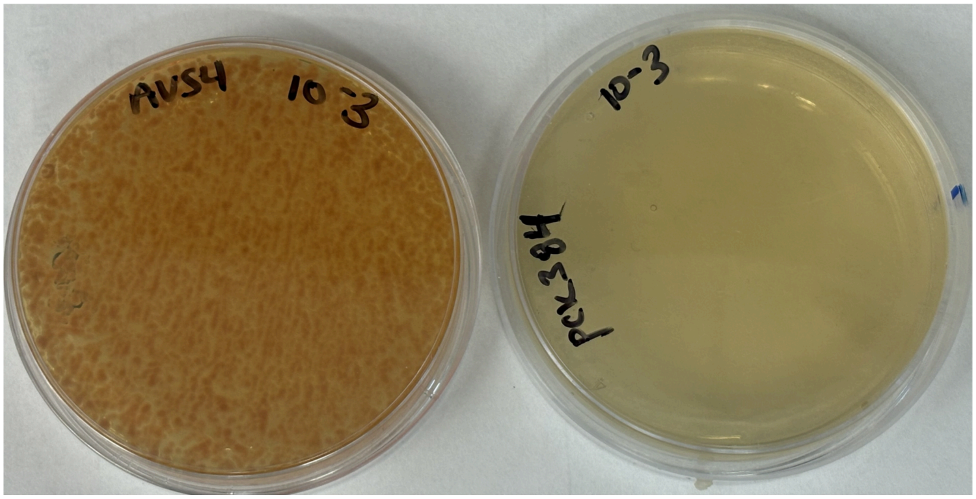


**Supplementary Figure 2.** Tri-parental conjugation was performed using the mini-Tn7 system, pJMP1039 (R6K-based Tn7 helper plasmid) and pCK383 (attTn7 cargo plasmid carrying an mRFP reporter and spectinomycin resistance marker), with *R. sphaeroides* as the recipient. (**left**) Viable recipient cell density was determined by serially diluting the same OD-normalized recipient culture used for conjugation (10-fold dilutions) and plating onto non-selective medium. Based on this recipient count and the absence of any detected transconjugants, the calculated detection limit for this experiment was $\leq 10^{-10}$ transconjugants per recipient CFU. (**right**) Representative selective plate (spectinomycin, $10^{-3}$ dilution). No transconjugant colonies were recovered at any dilution across the full plated series ($10^{-1}$ to $10^{-7}$).

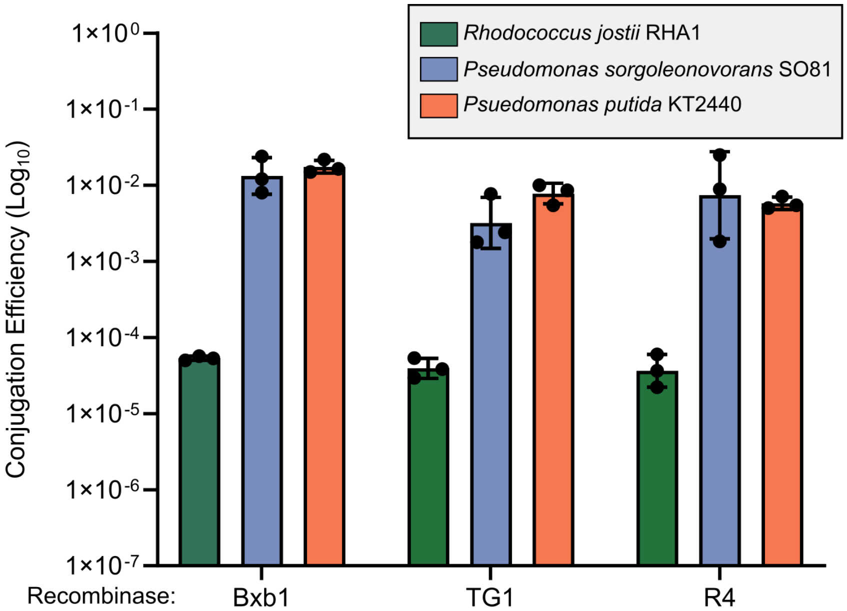


| | Bxb1 | TG1 | R4 |
|---|---|---|---|
| RHA1 | 87% | ND | ND |
| SO81: | 70% | 70% | 43% |
| KT2440: | 100% | 65% | ND |

**Supplementary Figure 3.** Conjugation efficiencies and integration accuracies shown across *Rhodococcus jostii* RHA1, *Pseudomonas sorgoleonovorans* SO81, and *Pseudomonas putida* KT2440 for Bxb1, R4, and TG1 integrases. Conjugation efficiency is defined as the number of transconjugant colonies normalized to recipient colony-forming units (CFUs) and is plotted on a $\log_{10}$ scale. Each bar represents the mean of biological replicates ($n$ = 3), with individual data points shown. The table below summarizes integration accuracy, defined as the percentage of transconjugant colonies exhibiting correct, on-target integration as confirmed by colony PCR ($n$ = 23 colonies per integrase and species tested). ND indicates conditions where colony PCR failed, resulting in incomplete integration accuracy determination for the indicated recombinases.

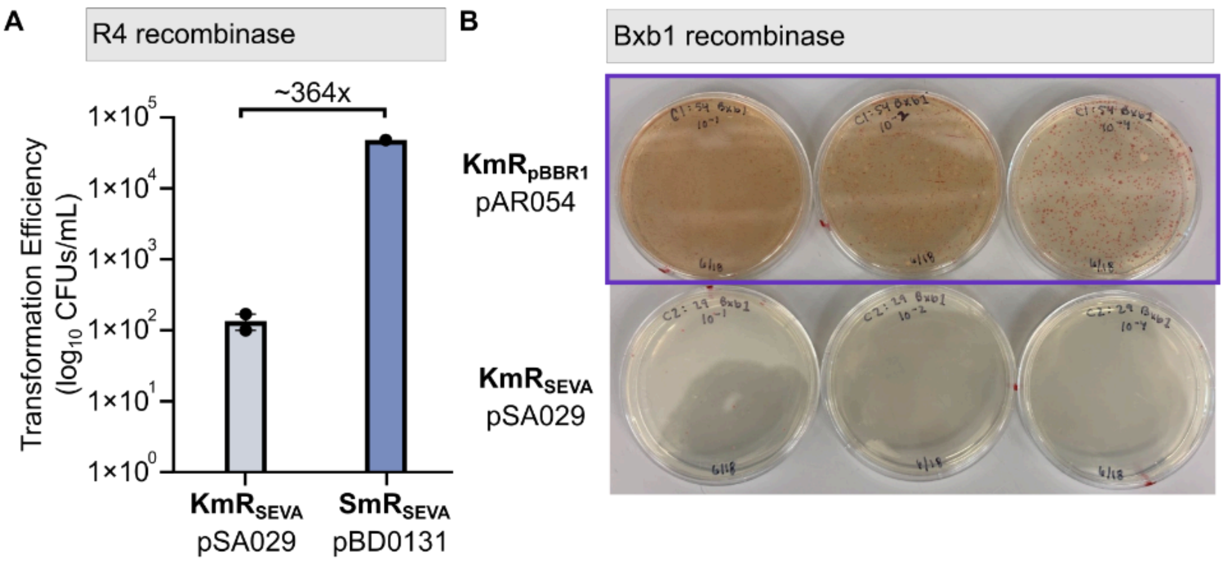


**Supplementary Figure 4.** Conjugation efficiencies in *R. sphaeroides* 2.4.1 using different antibiotic resistance markers. (A) Bar plot shows conjugation efficiency for two different payload/*attP* plasmids that differ in their resistance cassette: either a kanamycin resistance cassette ($Km^R$-SEVA; pSA029) or a spectinomycin ($Sm^R$-SEVA; pBD0131) resistance cassette. Both cassettes are from the SEVA collection. Data are representative of the average of replicate dilution plates (*n* = 3), where colonies were countable. (B) Representative images of conjugation plates following cSAGE integration with Bxb1 integrase and an *attP* plasmid containing either a $Km^R$ cassette cloned from the SEVA collection (pSA029) or a $Km^R$ cassette cloned from the replicating vector pBBR1MCS-2 (pAR054).

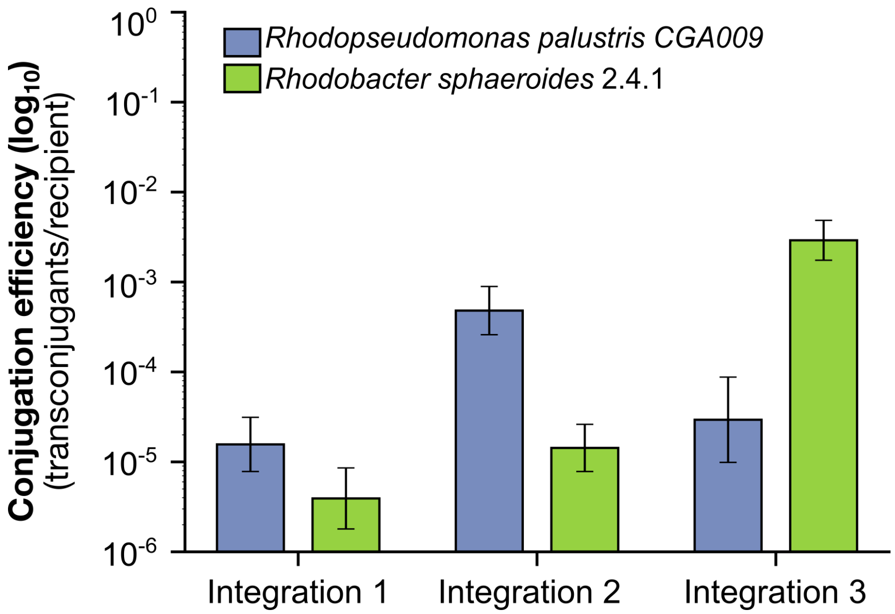


**Supplementary Figure 5.** Conjugation efficiencies after 3 sequential integrations in *Rhodopseudomonas palustris* CGA009 and *Rhodobacter sphaeroides* 2.4.1. Data plotted in order of integration: Round 1: R4 integrase + mTagBFP2/$Sm^R$; Round 2: Bxb1 integrase + mRFP/$Km^R$; Round 3: TG1 integrase + sfGFP/$Gm^R$. Control recipient-only plates were used to quantify conjugation efficiency as transconjugants per viable recipient. Individual data points represent conjugation efficiencies of replicate dilution plates ($n \geq 2$), with error bars showing standard deviation.

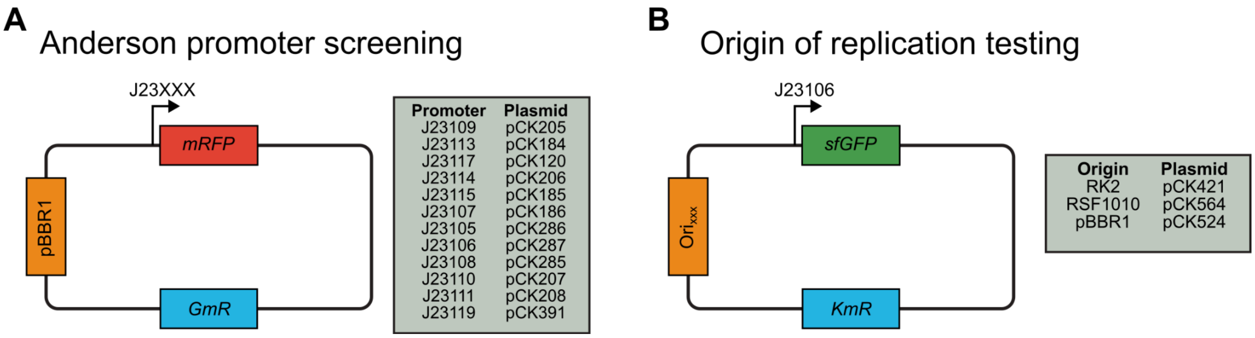


**Supplementary Figure 6.** Replicating vectors. (A) Library of Anderson promoters cloned upstream of mRFP in pBBR1MCS-2 backbone, used to benchmark promoter activity in *R. palustris* and *R. sphaeroides*. Each plasmid contains a gentamicin (GmR) resistance cassette. This plasmid series provides a resource for synthetic promoter characterization in PNSB (promoter sequences available at: https://parts.igem.org/Promoters/Catalog/Anderson). (B) Origins of replication tested in *R. sphaeroides*. Each plasmid contains an *sfGFP* under the J23106 promoter and a kanamycin (KmR) resistance cassette.

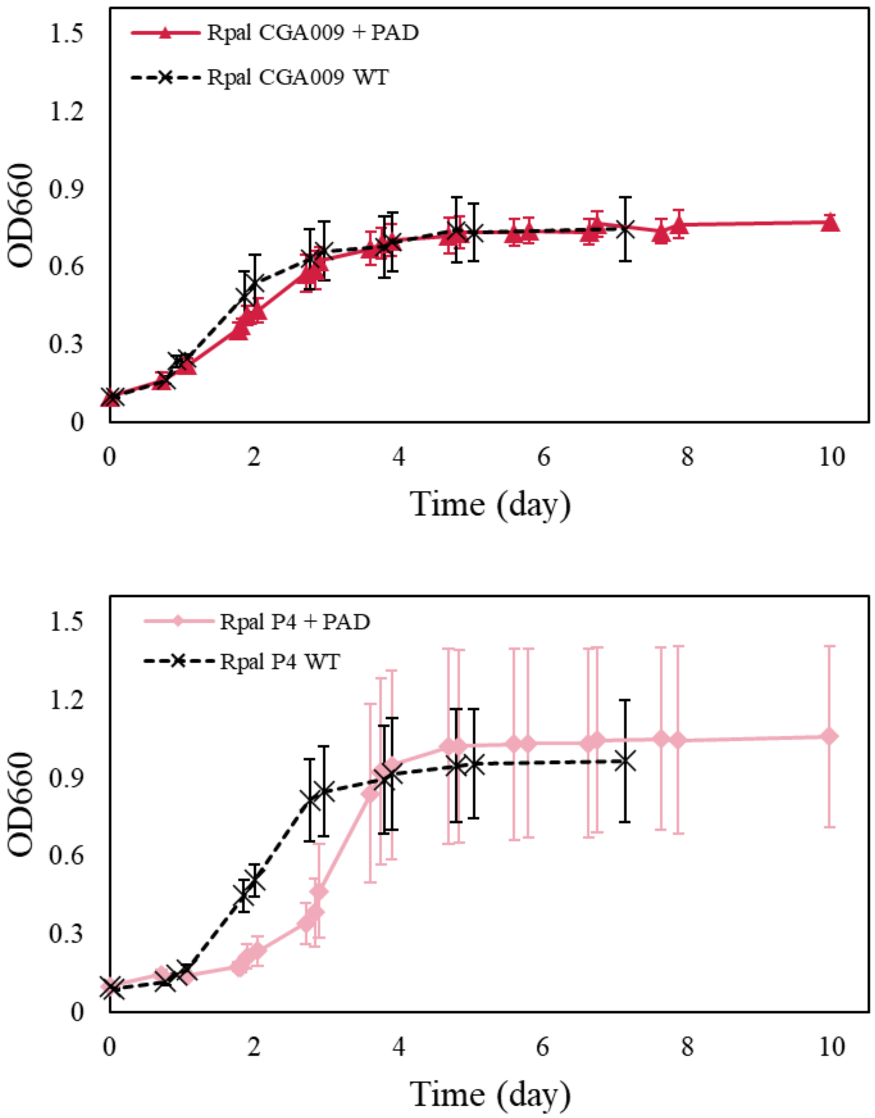


**Supplementary Figure 7.** Growth curves of engineered and wild-type *R. palustris* strains. Optical density at 660 nm ($OD_{660}$) was monitored over time for *R. palustris* CGA009 and P4 engineered strains expressing BaPAD or wild-type (WT) controls. Data points are the averages ± s.d. of three biological replicates.

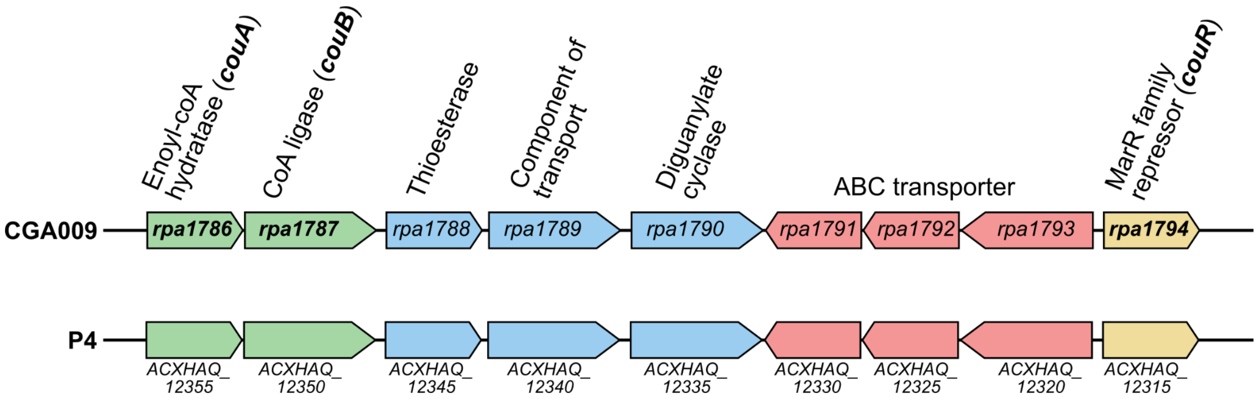


**Supplementary Figure 8.** Comparison of the *p*-coumarate catabolic pathway operon in *Rhodopseudomonas palustris* CGA009 (top) and *R. palustris* P4 (bottom). Gene content, order, and orientation are conserved between strains.

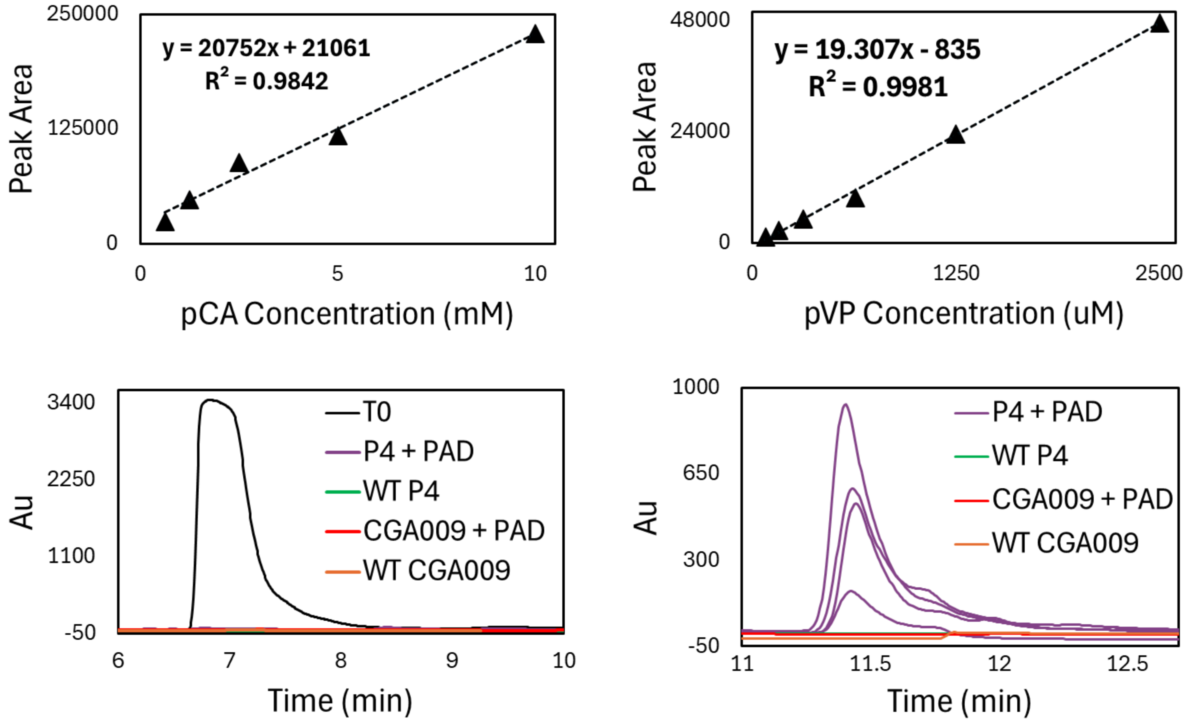


**Supplementary Figure 9.** Standard curves and representative chromatograms for *p*-vinylphenol (*p*-VP) and *p*-coumarate (*p*-CA) from *R. palustris* P4 and CGA009 bioproduction experiments. Calibration curves for *p*-CA (top left) and *p*-VP (top right) were generated from known concentrations and used to relate peak area to metabolite concentration via linear regression. Representative chromatograms show *p*-CA detection at ~7 min (bottom left) and *p*-VP detection at ~11.5 min (bottom right) for abiotic controls at the time of inoculation ($T_0$), BaPAD engineered strains (+PAD), and wild-type (WT) controls. The peak area was used to quantify metabolite concentrations in culture supernatants from bioproduction experiments according to known concentrations from the standard curves.

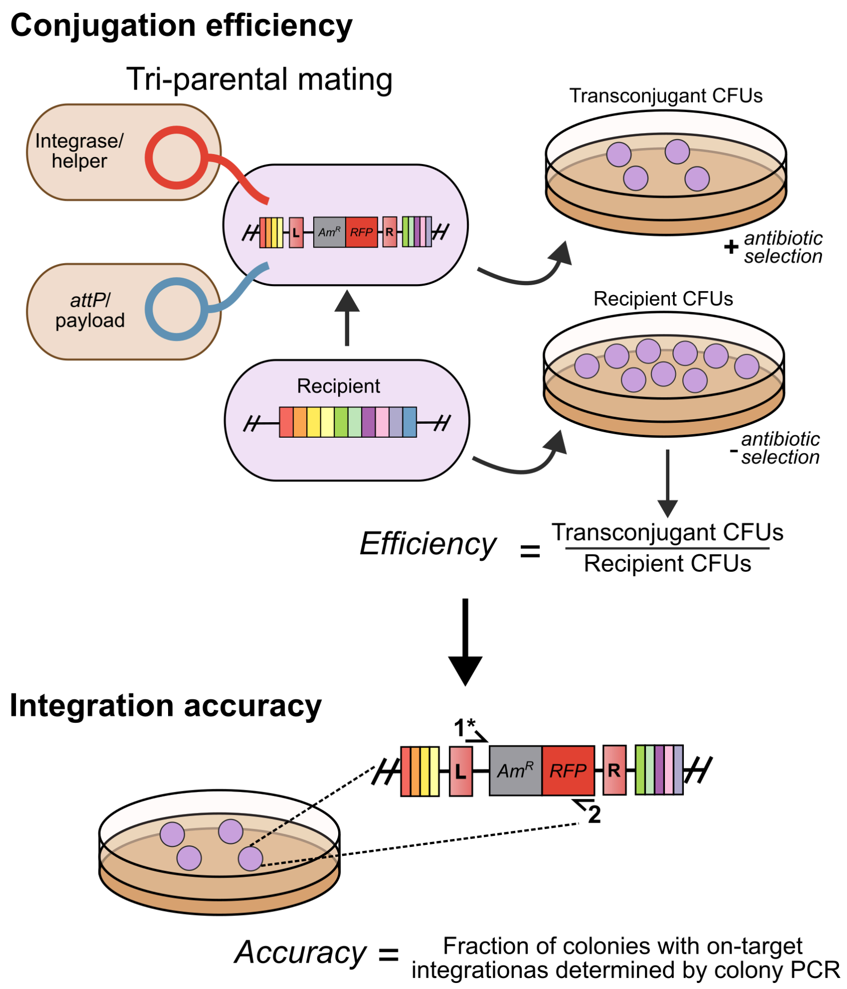


**Supplementary Figure 10.** Tri-parental conjugation assay used to assess conjugation efficiency and integration accuracy of cSAGE. (Top) Two *E. coli* donor cells, one carrying the integrase/helper plasmid and the other the *attP*/payload plasmid, are mated with a SAGE-enabled recipient strain by tri-parental conjugation. The mating mixture is plated on a selective medium lacking DAP to recover transconjugant CFUs. Recipient-only control plates, plated without selection, are used to determine total viable recipient CFUs. Conjugation efficiency is calculated as transconjugant CFUs divided by recipient CFUs. (Bottom) Integration accuracy is assessed by colony PCR using a forward primer flanking the targeted *attB* site and a reverse primer within the integrated cassette, confirming on-target integration at the intended site. Accuracy is calculated as the fraction of colonies yielding the expected on-target PCR product.

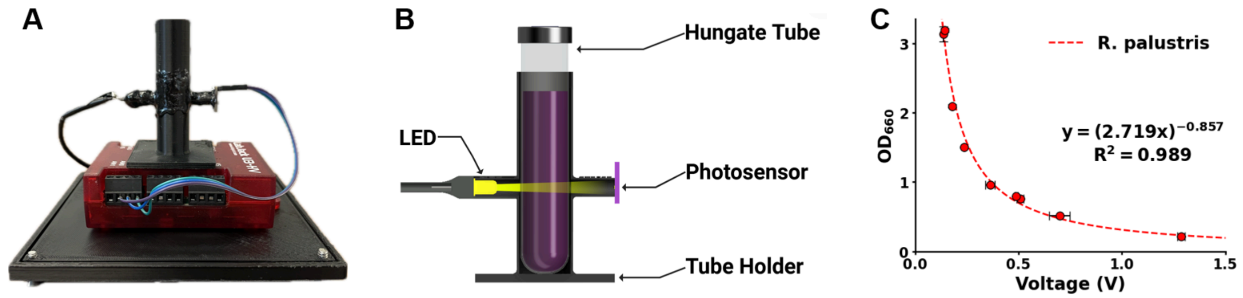


**Supplementary Figure 11.** Optical density sensor device and calibration curves for *Rhodopseudomonas palustris*. (A) Photograph of OD sensor device including sensor apparatus, LabJack U3-HV data acquisition device, and protective case. (B) Cross sectional diagram of OD sensor apparatus construction including 596 nm LED, TEMT6000 photosensor, and 3D-printed PLA tube holder. (C) Species specific $OD_{660}$ vs. voltage calibration curve for *R. palustris* generated using a benchtop spectrophotometer.
